# Full-Support Conditional Clade Distribution for Bayesian Tree Topology Testing

**DOI:** 10.64898/2025.12.16.694737

**Authors:** Zijing Yang, Jonathan Klawitter, Remco Bouckaert, Alexei J. Drummond

## Abstract

Bayesian phylogenetic inference uses Markov chain Monte Carlo sampling to estimate the posterior distribution of phylogenetic trees. However, the complex geometry of tree space makes these distributions difficult to characterise. Conditional clade distributions (CCDs) address this by providing a tractable model of posterior tree distributions, but existing parameterisations do not provide full support over tree space and exhibit complementary strengths. We introduce a new family of models, called *K-depth regularised conditional clade distributions (KRegCCD)*, which apply regularisation to balance sample fidelity against overfitting, thereby combining the strengths of existing parameterisations. KRegCCD achieves full support by extending the model to unexplored regions of tree space through a structured modelling of unseen clades. We show that KRegCCD outperforms existing models in posterior estimation while maintaining comparable point estimates. We further introduce an efficient cross-validation procedure for optimising regularisation parameters for individual posterior tree sets. Finally, we demonstrate that KRegCCD enables Bayesian tree topology testing by allowing candidate topologies to be evaluated against posterior tree distributions through their credible levels.

## 1 Introduction

Phylogenetics seeks to reconstruct the evolutionary history of organisms by analysing molecular sequence data. Bayesian inference [10, 32, 55, 75] has become widely adopted as a coherent probabilistic framework for estimating posterior distributions of trees and associated model parameters through Markov chain Monte Carlo (MCMC) sampling [28, 56]. Through repeated exploration of the parameter space, MCMC generates thousands of samples that approximate the true posterior distribution.

Among the parameters inferred in Bayesian phylogenetics, the tree topology is of particular importance, as it describes the evolutionary relationships for a set of taxa. Due to the high-dimensional and non-Euclidean nature of tree space [4, 6, 25], an MCMC sample cannot adequately represent the posterior distribution except in trivial cases. Continuous tree statistics such as the root age or the total tree length are routinely summarised by their full marginal distributions, but the topology, being discrete and high-dimensional, resists such treatment, and posterior samples of tree topologies are traditionally summarised by a single point estimate [29]. While such a summary provides a convenient visual and analytical reference, it inevitably discards much of the information contained in the posterior topology distribution, particularly regarding uncertainty in alternative topologies with substantial support. A finite sample also cannot assign a reliable probability to an individual topology. For any non-trivial analysis the number of topologies carrying appreciable posterior mass far exceeds any feasible sample size, so almost every distinct topology is visited at most once, and its Monte Carlo frequency, either 1*/N* or zero, is uninformative about its true posterior probability. Only low-dimensional summaries of the sample, such as clade and clade-split frequencies, are reliably estimable, which motivates modelling the posterior through such summaries rather than through raw topology counts. This has motivated the development of probabilistic representations that approximate the entire posterior tree distribution, rather than reducing it to a single representative tree.

Several models have been developed to address the challenges posed by the complexity of tree space [3,5,11,31,34,41,77]. Among these, the *conditional clade distribution (CCD)* approach is particularly effective, as it enables efficient and tractable approximation of posterior tree distributions. Introduced by Höhna and Drummond [31] and refined by Larget [41], a CCD provides normalised probabilities for all represented trees, and enables direct sampling from the distribution. CCD factorises the probability of a tree into conditional probabilities over recursively defined clade splits. This allows both calculating tree probabilities and sampling to be performed efficiently despite the large size of tree space. It has been applied in various contexts, including measuring phylogenetic entropy [42], detecting conflict among data partitions [42], finding rogue taxa [37], estimating credible levels of trees [38], species tree - gene tree reconciliation [69] and guiding tree proposals for MCMC runs [31].

One parameterisation, CCD0, models the posterior by per-clade parameters. It covers all trees that can be built by observed clades. This approach has been shown to produce more accurate point estimates than traditional methods such as the maximum clade credibility tree [5]. However, for some datasets it provides a poor approximation of the full posterior distribution [38]. Another parameterisation, CCD1 [31,41], instead models the conditional probability of each observed split. This representation better captures the shape of the posterior distribution, though its point estimates are less accurate than those of CCD0 [5]. One of our goals is to combine these complementary strengths into a single model.

Existing CCD parameterisations fundamentally rely on the finite set of clades and clade splits observed in the MCMC sample. Their probabilities are estimated directly from observed frequencies, so any clade or clade split that is never encountered is assigned zero probability. Zero probabilities arise in two ways: unobserved clade splits cannot occur within observed clades, and unobserved clades cannot be generated at all. This limitation is particularly important because finite MCMC samples represent only a tiny fraction of the vast tree space, especially for large datasets. As a result, standard CCDs model only the observed portion of the posterior distribution, and therefore do not provide full support over tree space.

To address these dual limitations, we introduce *K-depth regularised conditional clade distributions (KRegCCD)*, a regularisation framework that provides full-support probability estimates over tree space through a two-stage approach. First, we extend CCD1 by incorporating additional clade splits from CCD0 and applying pseudocount regularisation (a form of additive smoothing), enabling the mixture of the two models. In contrast to Zhang and Matsen [77], who also use pseudocounts but maintain fixed parameters and scale counts strictly by observed frequencies (leaving unobserved splits with zero probability), this step explicitly expands the set of candidate splits and optimises the pseudocount values. Second, to ensure support across the whole tree space, we reserve an escape probability for unobserved clades. Specifically, at a clade *C*, a small reserved probability mass (the clade’s escape probability) allows the model to “escape” the set of MCMC-observed clades and splits and assign probability to previously unobserved clades. To this end, we approximate the number of unobserved clades below *C* up to a depth *K*, giving rise to the name; details explained below. Together, these two stages define a tractable tree distribution that both improves posterior estimation and assigns non-zero probability to every tree topology in tree space.

An application enabled by KRegCCD is tree topology testing: assessing whether a topology is supported by sequence data, or can be rejected in favour of alternatives. This is a longstanding problem in phylogenetic inference, and classical approaches frame it as comparison of candidate trees by their likelihoods, asking whether the difference between them is statistically significant. Early approaches include the Kishino-Hasegawa test [35], which compares the log-likelihoods of two a priori specified trees using bootstrap sampling, and the Shimodaira–Hasegawa test [63], which extends this framework to multiple trees selected a posteriori, including the maximum-likelihood tree, while accounting for selection bias. The approximately unbiased test [62] improved bias correction through multiscale bootstrap resampling. The Swofford-Olsen-Waddell-Hillis test [68] takes a parametric bootstrap approach, evaluating whether one topology is significantly better than an alternative topology. Strimmer and Rambaut [67] introduced expected likelihood weights to quantify relative support among candidate trees and construct confidence sets based on cumulative likelihood weights.

While these methods provide rigorous statistical tools for testing tree topologies, they generally operate on a finite set of explicitly specified candidate trees and require likelihood calculations for each topology under consideration. Their conclusions are therefore conditional on the selected set of topologies. Furthermore, they assess topologies individually based on likelihood support and do not exploit the shared clade structure between related trees. KRegCCD, on the other hand, approximates the posterior distribution over the entire tree space from a finite posterior sample by modelling clade structure, providing a probability distribution that can be queried for arbitrary candidate trees. Specifically, the credible level of a candidate tree under a KRegCCD constructed from another posterior provides a Bayesian measure of its support in that posterior.

The contributions of this paper are as follows. We introduce regularisation steps for CCDs: split regularisation, which smooths the conditional clade probabilities over an expanded set of splits, and clade regularisation, which assigns probability to previously unseen clades. We combine these into KRegCCD, a family of CCD models that integrates the strengths of CCD0 and CCD1 while providing full support over tree space. We demonstrate that KRegCCD estimates the posterior tree distribution more accurately than both CCD0 and CCD1, while matching or improving on the point estimates of CCD1. To assess this, we propose a cross-validation procedure that evaluates how well a CCD model predicts held-out posterior trees. We give an efficient procedure for optimising the regularisation parameters, and evaluate it on simulated and real data. Finally, we show how KRegCCD enables Bayesian tree topology testing, allowing arbitrary candidate topologies to be evaluated against a posterior distribution through their credible levels.

## 2 Methods

In this section, we first introduce the basic notation and definitions required for this paper. Building on a general overview of the CCD framework, we then introduce KRegCCD with split and clade regularisation. Finally, we detail the datasets and experimental setup.

### 2.1 Phylogenetic Trees

Let *X* be a set of *n* taxa, say, *X* = {1, …, *n*}. A *rooted phylogenetic topology T* is a rooted tree with its leaves bijectively labelled with *X*. Let *V* (*T*) be the vertex set of *T* and let *E*(*T*) be the edge set of *T*. We call *T binary* if each interior vertex of *T* has exactly two children (i.e., out-degree two). A partial order on *V* (*T*) is established as *u* ≤ _*T*_ *v* if there is a path from the root to *u* that passes through *v*. In such a case and if *u ≠v*, we say that *v* is an *ancestor* of *u* and *u* is a *descendant* of *v*.

Let *T* be a phylogenetic tree on *X* and *v* a vertex of *T*. Let *T* (*v*) denote the subtree of *T* rooted at *v*. A subset *C* ⊆ *X* (potentially *C* = *X*) is a *clade* of *T* if there exists a vertex *v* ∈ *V* (*T*) such that the leaves of *T* (*v*) correspond exactly to the elements of *C*. Equivalently, a clade *C* is a subset of *X* such that all taxa in *C* share a common ancestor in *T*, and no taxon outside *C* shares that ancestor. A *clade split S* = {*C*_1_, *C*_2_} of a clade *C* is a pair of clades *C*_1_ and *C*_2_ with *C*_1_ ∪ *C*_2_ = *C* and *C*_1_ ∩ *C*_2_ = ∅. We also use the notation *C*_1_ ∥ *C*_2_ for *S* and *π*(*S*) to denote the parent clade *C* of *S*.

#### Remark.

Throughout, we now use the term “tree” to refer to a tree topology and assume that all trees are rooted and binary.

### 2.2 Conditional Clade Distributions

A CCD is a tractable tree distribution that parametrises tree space and offers an advanced estimation of a posterior distribution over phylogenetic trees [5,31,41]. A CCD *D* is represented by a *CCD graph G*, which is a directed bipartite graph: its vertices fall into two disjoint classes, the clades *C* (*G*) and the clade splits *S* (*G*), and every edge joins a vertex of one class to a vertex of the other. More precisely, each clade is connected to all its clade splits; each clade split is connected to the clades that constitute it. The root of *G* represents the clade encompassing the entire taxon set *X* and the leaves of *G* are the individual taxa in *X*. Trees *represented* by (or *contained* in) a CCD (graph) are obtained by starting at the root and recursively selecting a clade split for each clade.

A CCD can be parameterised by clade split probabilities. For a clade split *S* of a clade *C*, the *conditional clade probability (CCP)* Pr_*π*_(*S*) of *S*, is determined empirically as the fraction of trees in the posterior sample that contain *S* among those containing clade *C*. Formally, with *f* denoting the frequency of a clade or clade split appearing in a set of MCMC sampled trees *T*, we have:

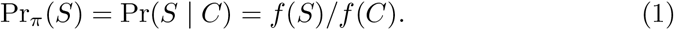

Note that this introduces a distribution on the clade splits of *C*. This parameterisation is known as *CCD1* [31, 41]. The probability of a tree *T* in a CCD1 distribution *D* is then given by the product of CCPs:

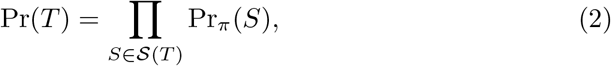

where *S* (*T*) is the set of clade splits of *T*. As shown by Larget [41], a CCD1 defines a probability distribution over trees, meaning that the probabilities of all trees in *D* sum to one. Viewed as a generative model, a CCD1 is a Bayesian network on the clade-split graph. A tree is generated top-down from the root clade, and the split chosen at a clade is conditionally independent of the rest of the tree given that clade. CCD1 is in this sense a first-order model: it conditions each split only on its immediate parent clade. This is a special case of the subsplit Bayesian networks of Zhang and Matsen [77], which condition on larger contexts. For example, Fig. 1 shows a posterior sample containing two topologies. The corresponding CCD1 graph represents two additional topologies that do not appear in the sample.

**Fig. 1:**
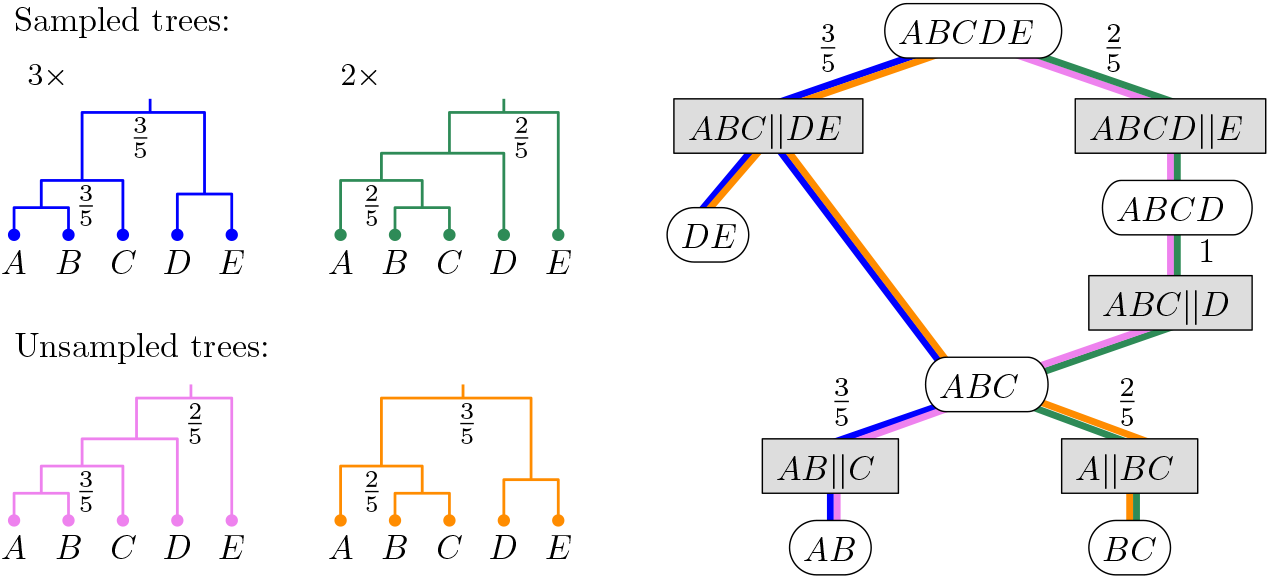
Posterior tree samples, the corresponding CCD1 graph built from the samples, and the unsampled trees it displays. The posterior sample contains five trees with two distinct topologies, sampled three and two times, respectively. Only clades *ABCDE* and *ABC* are split in different ways. The truncated CCD1 graph (cherry splits and leaf clades omitted) is annotated with clade split probabilities, estimated from the sampled trees. The probabilities assigned to the two sampled tree topologies in the CCD1 are 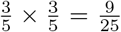 and 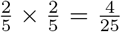, respectively. The CCD1 also assigns probability 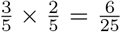 to each of the two unsampled trees. **Alt text:** A diagram shows sampled and unsampled trees alongside their corresponding CCD1 graph, with clade split probabilities annotated.

The *CCD-MAP tree* is the tree with maximum probability Pr(*T*) in the CCD, which can be computed efficiently using the dynamic program by Berling et al. [5].

The material in this subsection recapitulates the CCD framework of Larget [41] and Berling et al. [5] and is included only to establish notation; everything from Section 2.3 onwards is new to this paper. For further details and examples we refer to those works.

### 2.3 Regularised CCD

Standard CCDs assign zero probability to any tree containing an unobserved clade or split. This limitation arises from two sources. First, conditional clade probabilities are estimated directly from observed split frequencies, so unobserved splits receive zero probability. Second, the CCD graph itself contains only observed clades, preventing the generation of trees containing novel clades. We address these limitations by introducing a regularised CCD, called *KRegCCD*, which uses two complementary regularisation steps. Split regularisation smooths the conditional clade probabilities within observed clades, while clade regularisation assigns probability mass to previously unseen clades, resulting in a full-support tree distribution. Both steps parallel classical smoothing of sparse categorical data [16, 44]: split regularisation adds pseudocounts to the splits obtainable from observed clades, and clade regularisation reserves a fixed escape mass for trees that leave the observed clade set, spreading it over novel-clade regions with a geometric penalty per novel clade. For convenience, we refer to the model with split regularisation but without clade regularisation as *regCCD*.

#### Split Regularisation

In the first step towards KRegCCD, we regularise the conditional clade probabilities within each observed clade. CCD1 estimates CCPs directly from observed frequencies, assigning zero probability to splits absent from the posterior sample. Since finite MCMC samples inevitably miss possible splits, these estimates can overfit the observed sample. We therefore expand the set of possible splits and apply additive smoothing to the resulting conditional clade probabilities.

Additive smoothing is a standard regularisation technique for multinomial distributions [16,43,44]. It adds a positive pseudocount *α* to every outcome before normalisation, assigning non-zero probability to splits that were not observed in the sample.

To apply additive smoothing to CCDs, we first expand the set of clade splits considered by CCD1. This expansion is based on the CCD0 model [5], which contains all splits that can be formed from the observed clades in the posterior sample. For any clades *C, C*_1_, *C*_2_ in C(*G*) with *C*_1_ ∪ *C*_2_ = *C* and *C*_1_ ∩ *C*_2_ = ∅, we include the split {*C*_1_, *C*_2_} in *S*(*G*) (see an example in Fig. 2). Thus, the expanded graph contains the splits observed in CCD1 while adding splits implied by combinations of observed clades. We then apply additive smoothing over this expanded set, adding *α* to the empirical frequency of each observed split and assigning *α* to each expanded split, whose initial frequency is zero.

**Fig. 2:**
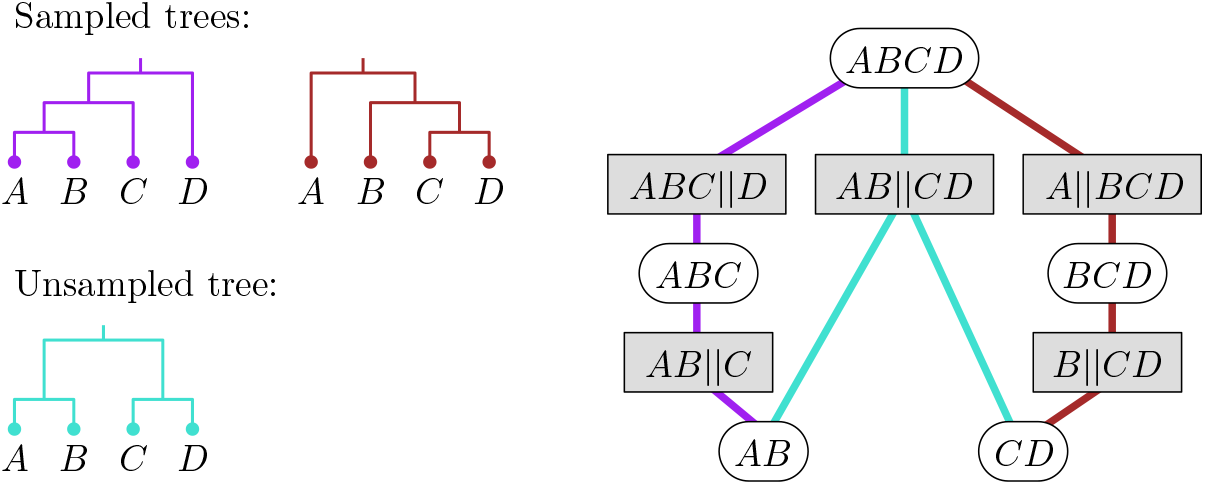
Posterior tree samples, the corresponding CCD0 graph built from the samples, and the unsampled tree it displays. CCD0 observes the clades *AB, CD* and *AB* ∥*CD*, and therefore contains the split *AB CD*. However, this split is absent from CCD1. **Alt text:** Tree samples and their CCD graphs illustrate how CCD0 can include a split absent from CCD1.

Let *D*_0_ denote the CCD0 and *D*_1_ the CCD1 built from the same tree set *T*. We construct a new CCD *D* using the graph of *D*_0_, which contains both the observed clade splits (*O* = *S*(*D*_1_)) and the expanded clade splits (*U* = *S*(*D*_0_) \ S(*D*_1_)). Recall that *f* (*S*) is the frequency (i.e. observed count) of *S* in *T* (and *f* (*C*) for a clade *C*). Note that for *S* ∈ *U*, we have *f* (*S*) = 0, and *f* (*C*) = ∑ _*S*∈*S*(*C*)_ *f* (*S*). For some *α* > 0, set the CCP of *S* with parent clade *C* in *D* as smoothed probability *θ*_*α*_(*S*) defined as:

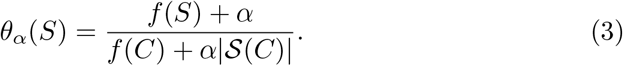

From a Bayesian perspective, this corresponds to using a Dirichlet(*α, α*, …, *α*) prior over the set of all observed and expanded clade splits and modelling the observed splits with a multinomial distribution.

The choice of *α* determines the strength of regularisation. The most common choice is *α* = 1, known as Laplace smoothing [44]. A more general approach, called Lidstone smoothing [43], uses 0 < *α* < 1 (often *α* = 0.5) which provides a more moderate adjustment. Note that *α* = 0 gives us the standard CCPs used by CCD1. We present a method for optimising *α* for a given dataset in Section 2.4. In addition to the standard pseudocounts 0 < *α* ≤ 1, we also tested an alternative pseudocount scheme that incorporates weights based on the prior probability of each clade split, so that clade splits receive non-uniform contributions rather than identical weights. We denote this case as *α* = prior.

### Clade Regularisation

Split regularisation assigns non-zero probability to unobserved splits, but only within the set of observed clades. Trees containing entirely new clades still receive zero probability, since those clades are absent from the CCD graph. The resulting distribution therefore still lacks full support over tree space. To overcome this limitation, we introduce clade regularisation, which assigns non-zero probability to trees outside regCCD.

We call clades and splits represented in the graph of a regCCD the *backbone* of the CCD. Splits in the backbone are assigned probabilities according to the *α*-smoothed CCPs in Eq. (3). We say a clade or split is *observed* if it belongs to the backbone, and *novel* otherwise.

A tree may contain clades or splits that are not represented in the backbone. We refer to a maximal connected region of a tree consisting of novel clades and splits as an *escape region*. For a tree *T*, we colour each clade or split red if it is observed and blue otherwise; Fig. 3 shows an example. Every escape region *B* has a parent clade *C*_*B*_, which is the smallest observed clade that contains the escape region. The escape region *B* must reconnect to the backbone, either through an observed internal clade or, eventually, through the singleton taxa, which are always observed. The downward *boundary*, denoted by *∂B*, of an escape region *B* is the set of observed clades at which the escape region reconnects to the backbone. The clades in *∂B* collectively partition the taxa of *C*_*B*_. We define the *depth i* of an escape region as the number of novel clades introduced between *C*_*B*_ and the boundary. If *B* contains *i* novel clades (i.e., has depth *i*), then |*∂B*| = *i* + 2. For example, in Fig. 3, the escape region contains one novel clade and therefore has three observed boundary clades.

**Fig. 3:**
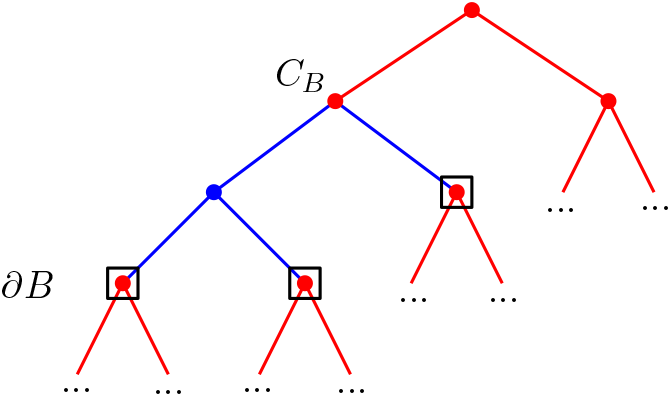
An illustrative section of a tree. The red regions represent observed clades and splits. The blue region is an escape region with parent *C*_*B*_, one novel clade, and three boundary clades *∂B*, marked by square boxes. **Alt text:** A tree diagram highlights an escape region within the surrounding observed tree structure, with parent clade and boundary clades.

Consider an observed clade *C* in the CCD backbone. An *escape subgraph* of the CCD below clade *C* is a connected subgraph that begins at *C*, contains only novel clades and splits internally, and reconnects to the backbone through a set of observed boundary clades. An escape subgraph of a CCD and an escape region in a tree share some underlying structure. Both are associated with a parent clade, contain novel clades, reconnect to boundary clades, and can be characterised by their depth. We apply the concepts of parent, boundary, and depth for escape regions to escape subgraphs, with the corresponding definitions adapted to the CCD graph. An example is shown in Fig. 4, where the red part represents the CCD backbone and multiple escape subgraphs are represented in blue. A detailed explanation of this example is provided in the large example at the end of Section 2.3.

**Fig. 4:**
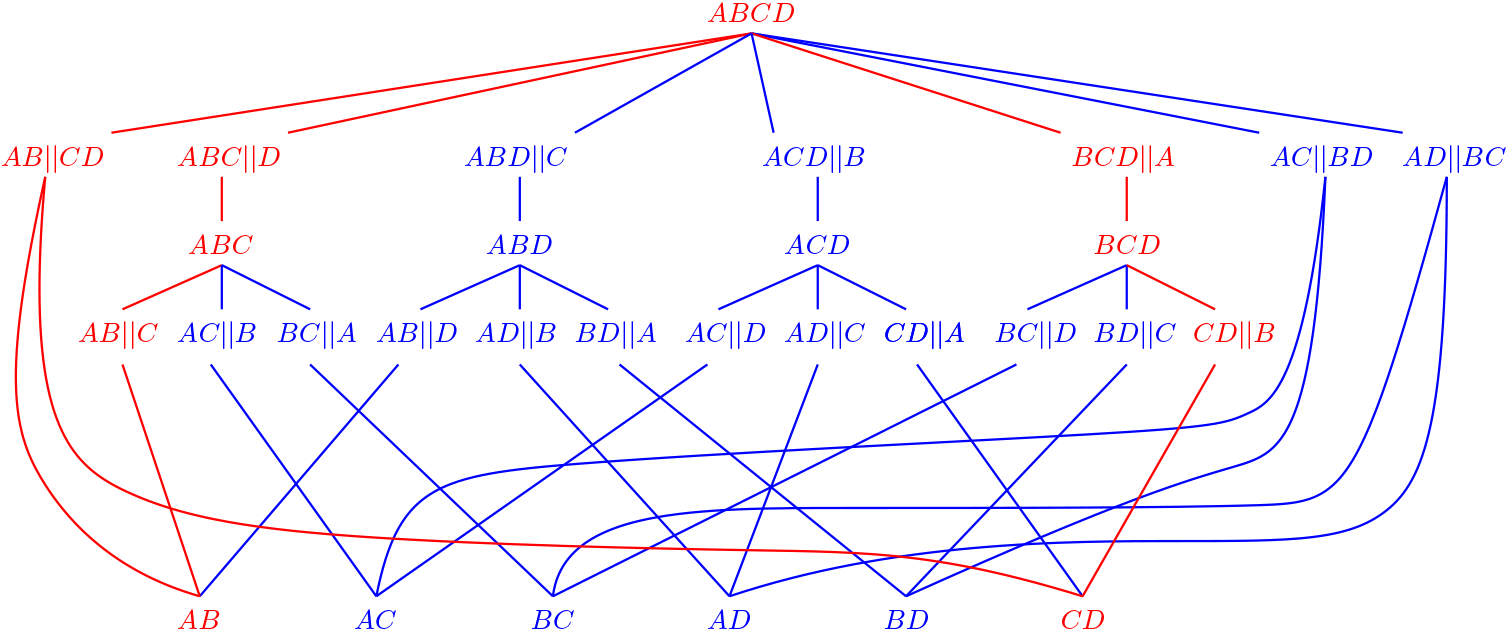
The KRegCCD graph over the full tree space of four taxa. The red region is the CCD backbone, which contains clades and splits that are either observed in the posterior sample or introduced by split expansion. The blue region consists of novel clades and splits that form multiple escape subgraphs (shown in Fig. 5). Singletons always belong to the red region and are omitted. **Alt text:** A four-taxon KRegCCD graph, with the backbone and several escape subgraphs representing previously unsupported parts of tree space.

**Fig. 5:**
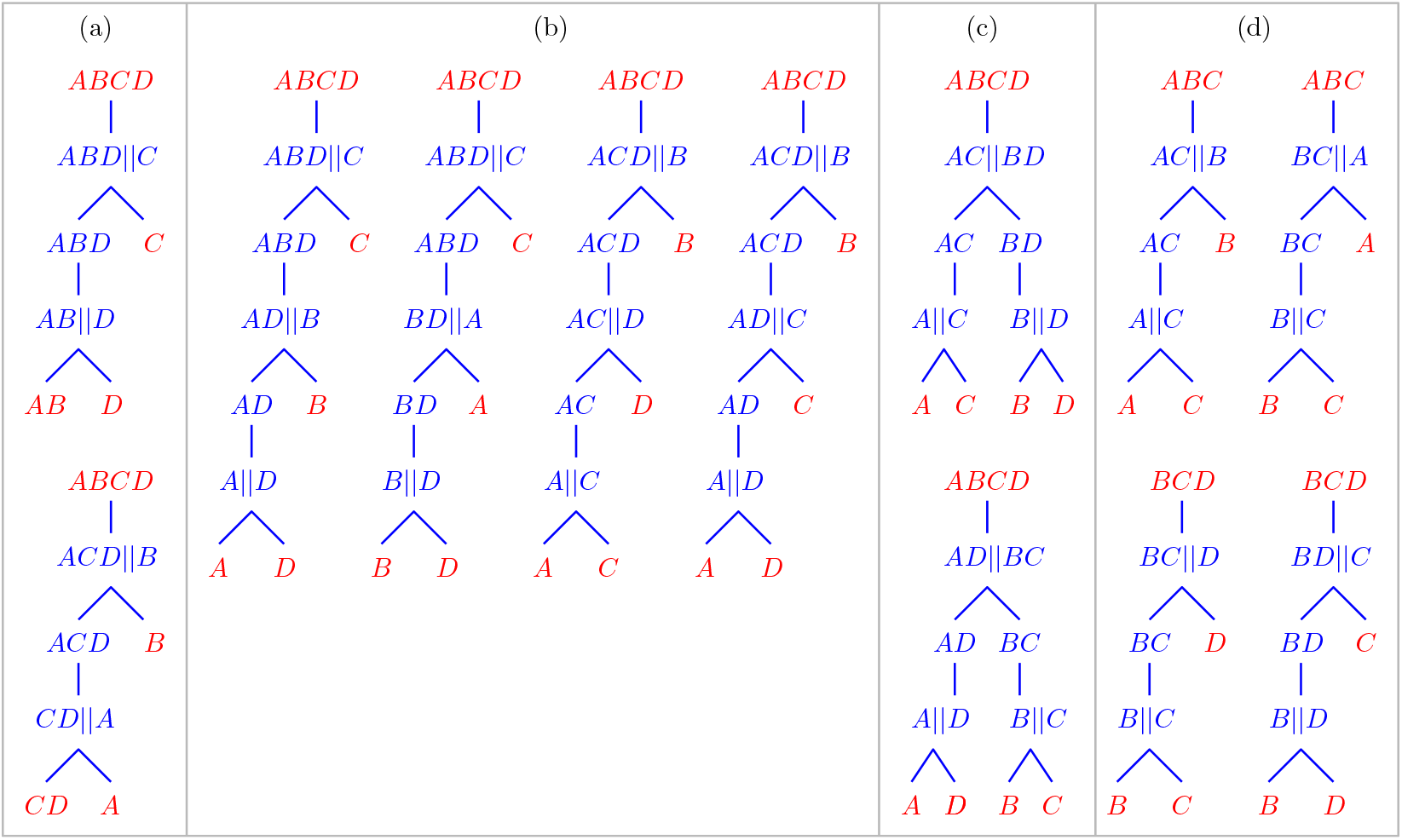
Escape subgraphs of Fig. 4. Red nodes are either parent or boundary nodes that are observed and therefore belong to the backbone. Blue regions consist of novel clades and splits. The escape subgraphs are grouped by parent, depth, and structure. (a) parent *ABCD*, depth 1; (b) parent *ABCD*, depth 2, chain structure; (c) parent *ABCD*, depth 2, sister-clade structure; (d) parent *ABC* or *BCD*, depth 1. **Alt text:** Four groups of escape subgraphs illustrate different combinations of parent clade, depth, and novel-clade structure within the four-taxon KRegCCD graph.

Let *N*_*i*_(*C*) denote the number of distinct escape subgraphs of depth *i* below *C*. Clade regularisation reserves a global escape probability *μ* at each observed clade *C*. At an observed clade *C*, a tree either continues along an observed split, with total probability 1 − *μ* distributed over all the observed splits of *C* according to *θ*_*α*_, or escapes into one of the ∑ _*i*≥1_ *N*_*i*_(*C*) possible escape subgraphs below *C*, with total probability *μ*. If all splits of *C* are in the backbone, then *μ* = 0 for *C*. The global escape probability *μ* remains the same whenever *μ ≠* 0. We set the default value of *μ* to 0.005 based on our experiments, while *μ* can be optimised by cross-validation for a specific dataset. Rather than distributing *μ* uniformly across escape subgraphs, we penalise deeper escape subgraphs geometrically. Let *ϵ*(*C*) be the clade-specific escape probability such that the total escape probability from *C* is equal to the global escape probability *μ*:

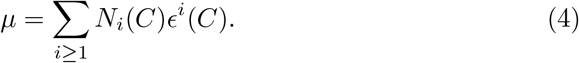

Since *N*_*i*_(*C*) depends on the structure below *C*, the resulting *ϵ*(*C*) is generally different for each clade. An example illustrating how *μ* is distributed is shown in Fig. 6 and explained in the large example at the end of Section 2.3.

**Fig. 6:**
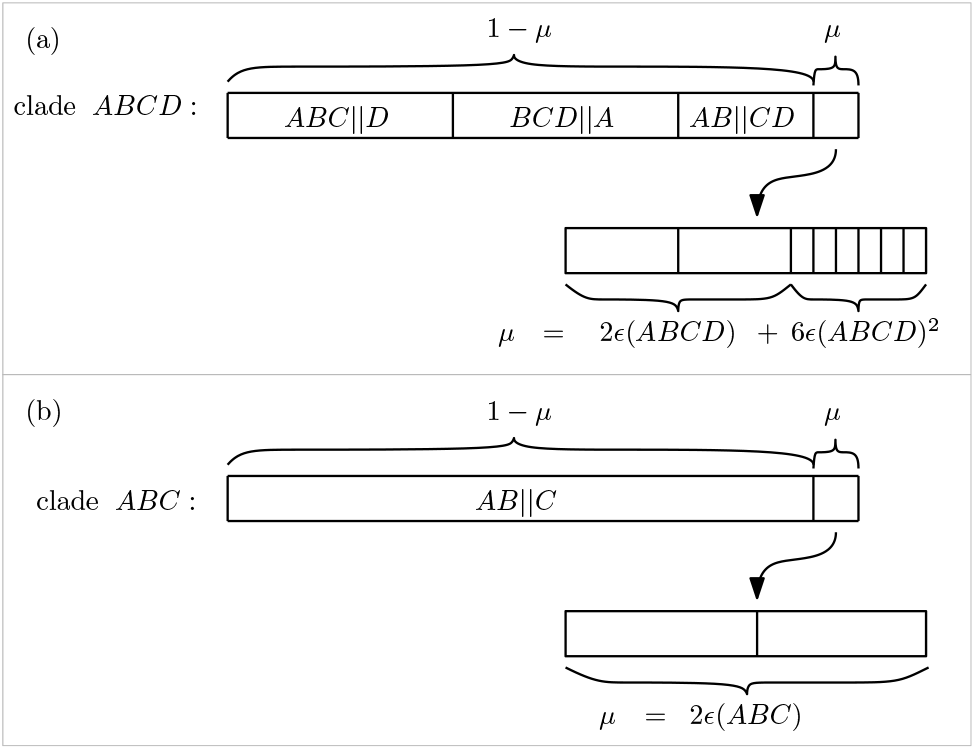
Probability allocation for clades *ABCD* and *ABC*. (a) Clade *ABCD* has three observed splits, which collectively receive probability 1− *α*. The remaining probability *μ* is distributed among the two depth-1 and six depth-2 escape subgraphs under *ABCD*, with the depth-1 subgraphs receiving a factor of *ϵ* each and the depth-2 subgraphs receiving a factor of *ϵ*^2^ each. (b) Clade *ABC* has one observed split, which receives probability 1 − *μ*. The remaining probability *μ* is distributed among the two depth-1 escape subgraphs under *ABC*, with each receiving a factor of *ϵ*. Clade *BCD* is treated similarly to clade *ABC*. **Alt text:** Two diagrams illustrate how probability is divided between observed splits and escape subgraphs at different depths within the KRegCCD model.

Computing the coefficients *N*_*i*_(*C*) exactly is #P-hard, as it amounts to counting set partitions over a non-laminar family of clades. In practice we therefore solve Eq. (4) using only the leading orders *i* ≤ 2, corresponding to a truncation depth of *k* = 2, while still scoring escape regions of any depth, so full support is retained. Since *ϵ*(*C*) is of order *μ/N*_1_(*C*), retaining orders *i* ≤ *k* leaves an omitted tail of order *O*(*μ*^*k*+1^): the escape probability actually assigned to a clade departs from *μ* by *O*(*μ*^3^) at *k* = 2. This is negligible at the values of *μ* selected by cross-validation (*μ* ≤ 0.05). We refer to this family of models as KRegCCD, where *k* is the truncation depth.

To evaluate the probability of a tree *T*, we decompose it into regions relative to the CCD graph. Let *R*(*T*) denote the set of observed splits in the backbone, and let *B*(*T*) denote the set of escape regions. For each escape region *B* ∈ *B*(*T*), let *C*_*B*_ be its parent clade and *i* its depth. The probability of *T* is therefore

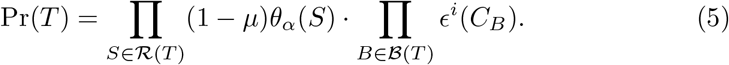

An example of calculating tree probabilities is shown in Fig. 7 and explained next.

**Fig. 7:**
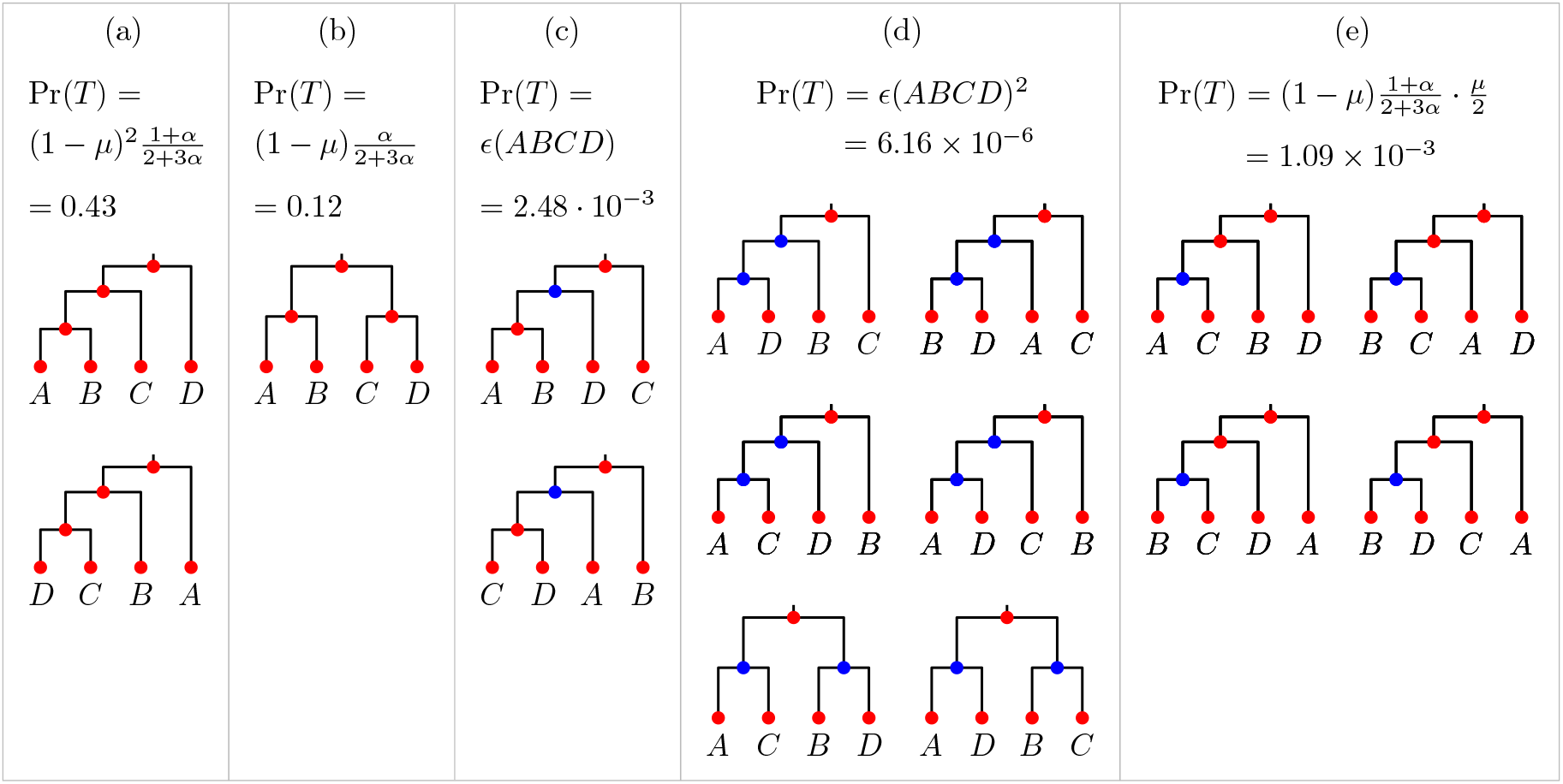
All 15 trees on four taxa, grouped into categories of trees with equal probability. The expression shown in each category denotes the probability of each individual tree in that category. The concrete probabilities assume *α* = 0.4 and *μ* = 0.005. Nodes are coloured by origin: red denotes clades observed in the posterior sample, while blue denotes novel clades. (a) Sampled trees. (b) Expanded tree. (c) Trees escaped from the root with 1 novel clade. (d) Trees escaped from the root with 2 novel clades. (e) Trees escaped from clade *ABC* or *BCD* with 1 novel clade. **Alt text:** All four-taxon trees and their probabilities, with red and blue nodes distinguishing observed and novel clades across five structural categories.

#### Example.

Suppose a posterior sample consists of two trees, each observed once: (((*A, B*), *C*), *D*) and (((*D, C*), *B*), *A*). The observed non-trivial clades are *ABC, AB, BCD*, and *CD*. The backbone is marked in red in Fig. 4. At the root clade *ABCD*, which is observed twice, there are three candidate splits: the two observed splits and *AB* ∥ *CD* obtained by split expansion. Their CCPs are calculated by additive-*α* smoothing:

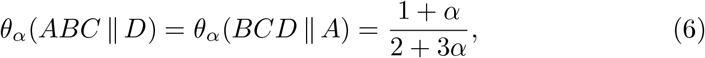

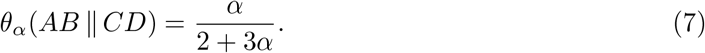

The two interior three-taxon clades each contain one split, so their CCP is 1. The escape subgraphs of Fig. 4 are marked in blue. For the root clade *ABCD*, there are two escape subgraphs introducing one novel clade (depth 1), shown in Fig. 5 (a); further there are six escape subgraphs introducing two novel clades (depth 2), four with the chain structure shown in Fig. 5 (b) and two with the sister-clade structure shown in Fig. 5 (c). This gives *N*_1_ = 2 and *N*_2_ = 6. Hence,

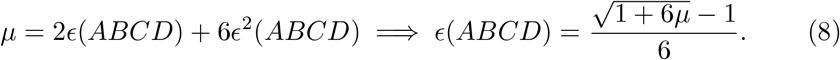

For clades *ABC* and *BCD*, there are two escape subgraphs for each clade, both introducing one novel clade (depth 1), shown in Fig. 5 (d). This gives *N*_1_ = 2, and

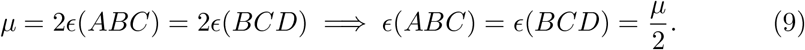

Fig. 6 shows how mu is distributed to these escape subgraphs. Each escape subgraph of depth *i* under a clade *C* receives a factor of *ϵ*^*i*^(*C*). More precisely, for clade *ABCD*, the two depth-1 escape subgraphs each receive *ϵ*(*ABCD*), and the six depth-2 escape subgraphs each receive *ϵ*^2^(*ABCD*). For clade *ABC*, the two depth-1 escape subgraphs each receive *ϵ*(*ABC*), and analogously for clade *BCD*.

The full tree space on four taxa contains 15 trees, all represented by KRegCCD. The probability of each tree is shown in Fig. 7. To obtain concrete probabilities, let’s assume that *α* = 0.4 and *μ* = 0.005.

Fig. 7 (a) The two trees contain only clades in the backbone, so their probability is (1 − *μ*)*θ*_*α*_(root split) multiplied by the CCPs of *AB* ∥ *C* and *CD* ∥ *B*, respectively. Since there are no expanded splits under clades *ABC* and *BCD*, the corresponding CCPs are each (1 − *μ*). Thus, each tree has probability 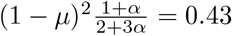.

Fig. 7 (b) The tree is covered by the backbone through split expansion. Its probability is (1−*μ*)*θ*_*α*_(*AB* ∥ *CD*) multiplied by the CCPs of the two cherry splits, both of which have CCP 1. This gives 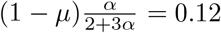.

Fig. 7 (c) Each tree contains one novel clade under the root and therefore has probability 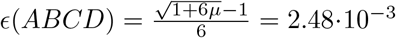, as derived in Eq. (8).

Fig. 7 (d) Each tree contains two novel clades under the root and therefore has probability 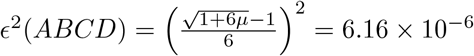.

Fig. 7 (e) Each of the trees have one novel clade under either clade *ABC* or *BCD*. So their probability is (1 − *μ*)*θ*_*α*_(root split) multiplied by *ϵ*(*ABC*) and *ϵ*(*BCD*), respectively. Each tree has probability 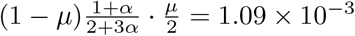, where 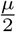 is derived in Eq. (9).

### 2.4 Model Evaluation

We evaluated KRegCCD using three complementary approaches: distributional accuracy (credible set coverage), predictive performance (cross-validation), and point estimation accuracy (distance to true tree). We first describe the datasets used, then discuss each evaluation approach in turn.

#### Datasets

We used well-calibrated simulation studies (WCSS) [47] datasets from Berling et al. [5], which were produced with LinguaPhylo [21] and BEAST2 [10] and grounded in simple models that have been extensively tested. Since these datasets are expected to pass a coverage analysis, they provide a reliable basis for CCD model assessment. The WCSS datasets are summarised briefly as follows:

Yule[*n*] For number of taxa *n* ∈ {50, 100, 200, 400}, we used 100 Yule tree [76] simulations (replicates). Each replicate has two independent MCMC runs, with the birth rate fixed at 25.0 (12.5 for *n* = 20), a HKY+G substitution model [27], the shape parameter for the gamma distribution of site rates using Lognormal(−1.0, 0.5), the transition/transversion rate ratio *κ* followed Lognormal(−1.0, 1.25), nucleotide base frequencies independently simulated for each replicate from Dirichlet(5.0, 5.0, 5.0, 5.0), sequence alignments of 300 sites, and the mutation rate fixed at 1.0.

Coal[*n*] For number of taxa *n* ∈ {40, 80, 160, 320}, we used 100 time-stamped coalescent [54] simulations, each with two independent MCMC runs, a population size *θ* drawn from Lognormal(−2.4276, 0.5), sequence alignments of 250 sites, the youngest leaf at age 0, the remaining leaf ages distributed uniformly at random between 0 and 0.2, and all other parameters as in the Yule simulations.

All replicates of Yule[*n*] and Coal[*n*] were confirmed to have converged, with excess burnin discarded.

In addition to the WCSS simulation datasets, we generated the RSV2 posterior as a controlled real-data case study to evaluate both posterior coverage and predictive performance under different regularisation strategies.

`RSV2` The `RSV2` strict-clock posterior tree set, containing 129 taxa, was obtained from an analysis of human respiratory syncytial virus subgroup A (RSVA) sequences based on the attachment glycoprotein (G) gene. The sequences were sampled from different geographic locations, with isolation dates spanning from 1956 to 2002 [78, 79].

We also analysed a large collection of real datasets from the literature:

`RealData` More than 100 posterior tree sets from thirty published studies [2, 7, 8, 12, 15, 17, 18, 23, 24, 26, 30, 33, 36, 39, 40, 45, 46, 48–50, 57–61, 64–66, 70, 72] that conducted Bayesian phylogenetic analyses, with the number of taxa ranging from 5 to 579, varied tree sample sizes, and model configurations. A burnin of 10% was applied to each tree set. A summary of these datasets is provided in Appendix S3.

We further used several datasets in RealData for Bayesian tree topology testing:

`Percidae` Two sets of 8,000 trees on 233 taxa from analyses with unconstrained topology priors and topology priors constrained by a higher-level taxonomic classification [2].

`Liocichla` Two sets of 40,000 trees on 6 taxa from analyses under codon-partitioned and gene-partitioned models [46].

`Rallidae` Two sets of 20,000 trees on 100 taxa from a combined five-marker shared-tree analysis and a mitochondrial-only tree analysis [66].

#### Credible Set Coverage

We evaluated the accuracy of posterior estimation by KRegCCD using credible set coverage. A *γ credible set C*_*γ*_ in Bayesian phylogenetics is the analogue of a credible interval for discrete parameters. It is defined as a smallest subset of elements that together account for at least *γ* of the posterior probability mass. The *credible level γ* is a value in the interval (0, 1], typically written as a percentage. For example, when *γ* = 0.1, the 10% credible set consists of the most probable trees, added in descending order of posterior probability, until their cumulative probability reaches 10%.

We used *probability-based credible set* introduced by Klawitter and Drummond [38] for our experiment, which works as follows. A probability-based credible set of a CCD *D* is based on probability thresholds for elements in *C*_*γ*_ obtained by sampling trees from *D*. Let *k* be the number of trees sampled from *D* (e.g. *k* = 10, 000). We get the probability of each sampled tree and sort them in decreasing order. Let *T*_1_, …, *T*_*k*_ be the sampled trees in this order. For each credible level *γ*_*i*_ ∈ {0.01, 0.02, …, 1.0}, let *j* = ⌈*γ*_*i*_ · *k*⌉ and *p*_*j*_ = Pr_*D*_(*T*_*j*_). Then *p*_*j*_ is set as the probability threshold for any tree to be contained in 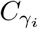. The credible level of a tree *T* is obtained by computing its probability Pr(*T*) in *D*, then finding the smallest *γ*_*i*_ such that 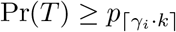.

For each replicate in `Yule[n]` and `Coal[n]`, we used the posterior tree sets from the two independent MCMC runs, assigning one as the *training set* and the other as the *testing set*. From the training set, we drew evenly spaced subsamples of sizes 100, 300, and 1000, and used each subsample to build CCD0, CCD1, and KRegCCD. For each CCD, we computed the probability-based credible sets. We then drew subsamples of the same sizes from the testing set and for each tree *T* in such a subsample, we computed the credible level of *T* in the corresponding CCD, counting the number of trees falling into each level (0.01, 0.02, …, 1.0).

### Cross-validation

We use cross-validation (CV) as a general framework for evaluating predictive performance and for optimising the regularisation parameters.

Given a posterior sample of *N* trees {*T*_1_, …, *T*_*N*_}, we partition the sample into *F* folds (subsets). For each fold *f*, we construct a CCD *D*^(−*f*)^ using the trees in the remaining folds, and compute the probability of each held-out tree under *D*^(−*f*)^. The predictive performance of a CCD is summarised by the mean log probability across all held-out trees:

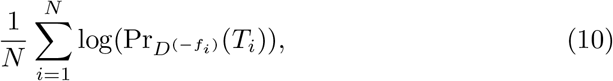

where *f*_*i*_ denotes the fold containing tree *T*_*i*_ and 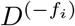 is the CCD constructed without that fold. Higher values indicate a more accurate approximation of the posterior distribution.

### Held-out predictive performance

To evaluate how well each model predicts unseen posterior trees, we divided each posterior tree set into two equal parts, or used two MCMC chains when available. The first part, or first chain, was used to train the CCD, and the second part, or second chain, was used as a held-out test set. For each tree in the test set, we calculated its probability under the CCD fitted to the training set. We then measured *predictive performance* as the mean log probability of the held-out trees. We also measured *support coverage*, defined as the proportion of held-out trees assigned non-zero probability by the model.

### Distance to Truth

To assess the effect of regularisation on point estimation, we measured the Robinson-Foulds (RF) distance between the CCD-MAP tree and the true tree. The *Robinson-Foulds (RF) distance* [53] is a metric quantifying the dissimilarity between two trees by counting the number of clades present in one tree but absent in the other. We used the relative RF distance, defined as the standard RF distance normalised by 2(*n* −2), where *n* is the number of taxa. We used subsamples of size 3, 10, 30, 100, 300, 1k and 3k to generate a CCD0, a CCD1, and a KRegCCD for all 100 simulations of each dataset Yule[n] and Coal[n]. The subsamples were evenly spaced across the original MCMC samples. From each CCD, we then obtained the MAP tree and measured its RF distance to the true tree.

## 3 Results

We first evaluate whether KRegCCD improves the estimation of posterior tree distributions and demonstrate the importance of structured regularisation. We then examine how regularisation affects point estimation, optimise the regularisation parameters, and finally demonstrate its application to Bayesian tree topology testing.

### 3.1 Tree Distribution

To evaluate the performance of KRegCCD in estimating the whole posterior distribution, we computed the fraction of trees from the testing set that fall within the credible sets constructed from the training set. The results for Yule100 are shown in Fig. 8, with similar results for the remaining Yule[n] and Coal[n] datasets provided in Appendix S1.

**Fig. 8:**
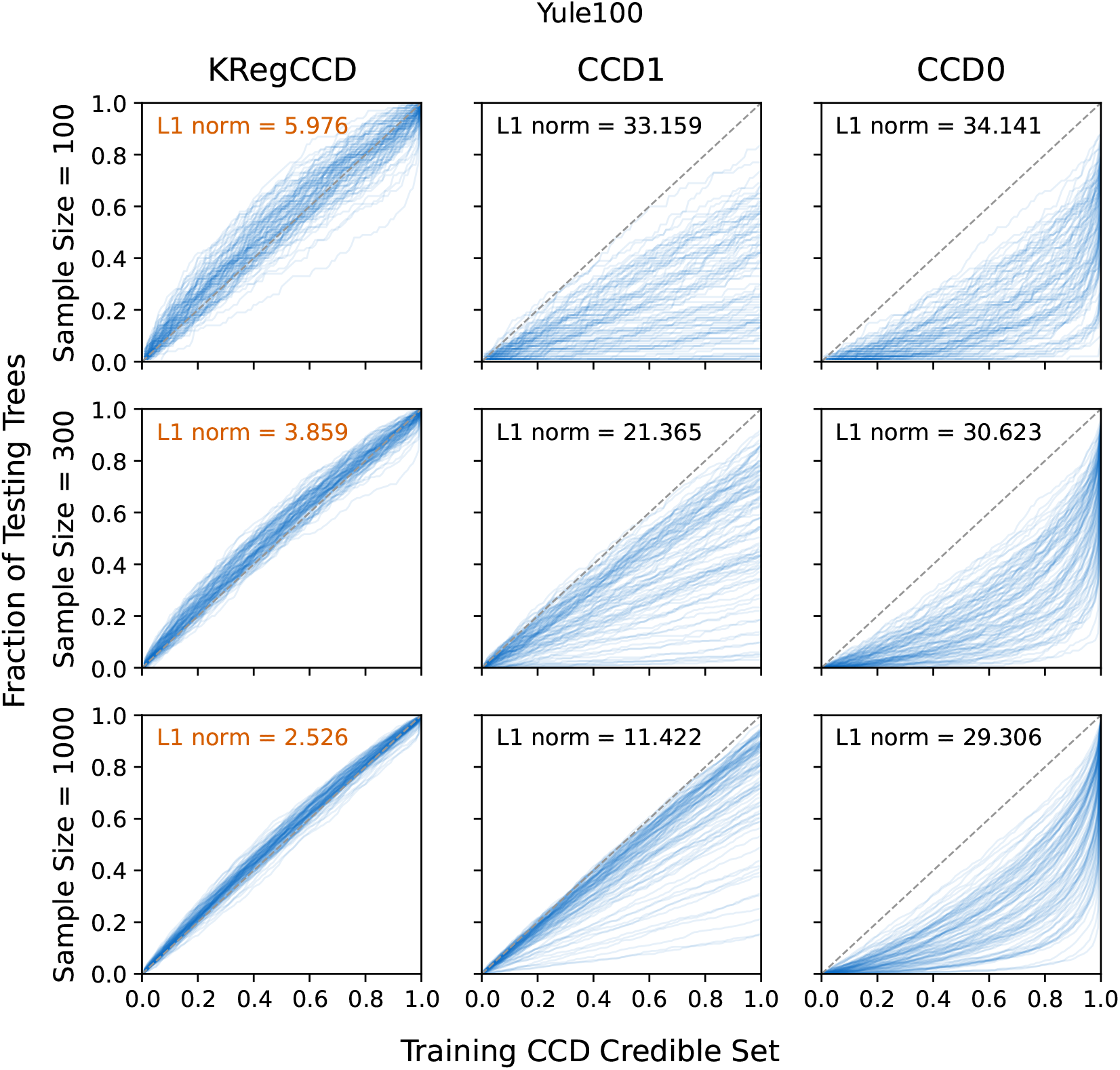
The accuracy of the full posterior estimation for Yule100, measured as the fraction of the testing trees in a range of CCD credible sets built from the training set and annotated with average L1 norms, with the lowest value for each sample size highlighted in orange. Models CCD0, CCD1, and KRegCCD with *α* = 0.4 and optimised *μ* are compared across sample sizes of 100, 300, and 1000. CCD0 performs the worst and largely underestimates posterior support, CCD1 provides better estimates than CCD0, and KRegCCD consistently achieves the highest accuracy. **Alt text:** A comparison of posterior estimation accuracy across sample sizes shows differences among KRegCCD, CCD1, and CCD0 using credible set coverage.

The first column of Fig. 8 represents KRegCCD, the second column CCD1, and the third CCD0. The rows correspond to different sample sizes. Along the discrete x-axis, the CCD credible sets for credible levels (0.01, 0.02, …, 1.0) are represented, and each y-axis shows the fraction of testing trees contained within the respective credible set. Each blue line represents a replicate, and the closer the lines are to the diagonal, the better the CCD estimates the posterior distribution. Each plot also shows the average L1 norms, which measure the distance of the blue line from the diagonal. For each sample size, the lowest norm is highlighted in orange.

If a line lies below the diagonal, the model assigns lower probability to test trees than warranted: the distribution is too diffuse, either spreading probability mass across too many trees or including spurious trees that dilute the probability of trees that actually occur in the posterior. Conversely, if a line lies above the diagonal, the model assigns higher probability to test trees than warranted: the distribution is too concentrated, overfitting to the sample.

In Fig. 8, CCD0 produced the loosest fit, with all lines lying far below the diagonal, CCD1 performed better than CCD0, but still largely underestimated tree probabilities. In contrast, KRegCCD produced substantially better fits.

Paired Wilcoxon signed-rank tests confirmed these improvements are highly significant (Table 1). KRegCCD achieved significantly lower L1 norms than both CCD1 and CCD0 across all datasets (*p* < 0.05 in all cases). For datasets with 50 or more taxa, KRegCCD outperformed CCD1 in 81–100 of 100 replicates, and outperformed CCD0 in 100 of 100 replicates for all datasets.

**Table 1:**
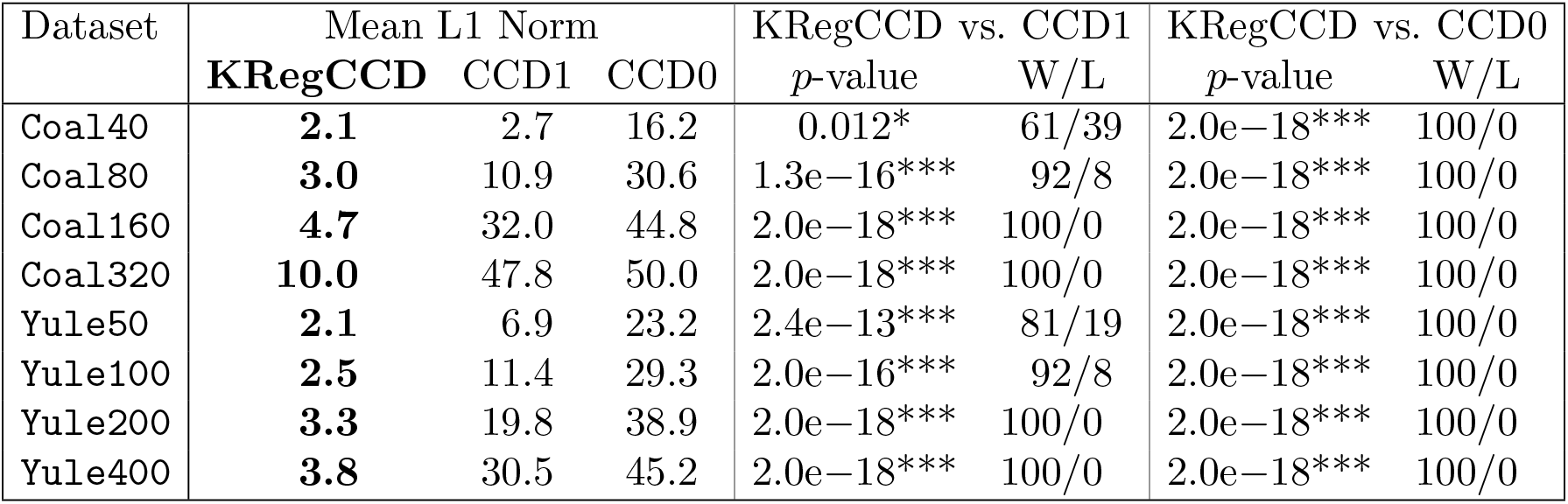
Statistical comparison of posterior distribution estimates (subsample size = 1000). Mean L1 norms (distance from ideal diagonal) and Wilcoxon signed-rank test p-values are shown. Lower L1 norm indicates better calibration. W/L (win/loss) denotes the number of replicates in which KRegCCD has a lower/higher L1 norm than the compared model. Significance: ^*^*p* < 0.05, ^**^*p* < 0.01, ^***^*p* < 0.001. P-values are not corrected for multiple testing; all results remain highly significant under Bonferroni correction.

The limited coverage of CCD0 and CCD1 arises because they do not assign probability to the entire tree space; consequently, some testing trees receive zero probability under the CCD constructed from the training sample. Therefore, even when considering the full credible set of the training CCD (credible level = 1.0), the fraction of testing trees included in the credible set may remain below 1. KRegCCD provides full support over the tree space, ensuring that all testing trees receive non-zero probability under the training model. As a result, the fraction of testing trees included in the full credible set (credible level = 1.0) always reaches 1.

### 3.2 Structured Regularisation

To demonstrate the importance of structured regularisation, we evaluated support coverage and held-out predictive performance, as described in Section 2.4, on several real datasets.

We compare four models. CCD1 is the unregularised baseline, using only splits sampled in the posterior tree set, and regCCD extends the set of allowed splits and applies *α*-smoothing; CCD1+*ϵ* gives the trivial way to achieve full support by using CCD1 as its backbone and reserving a small uniform escape probability *ϵ* for trees outside the backbone, whereas KRegCCD uses regCCD as its backbone and distributes an escape probability *μ* among trees outside the backbone according to the clade structure.

The coverage results are shown in Fig. 9 and the predictive performance results in Table 2. CCD1 and regCCD do not provide full support over the tree space. Across the 20 datasets, most have less than 100% coverage under both models, with only one dataset achieving full coverage under regCCD. Trees outside the model support receive zero probability, so their log probability is −∞. Full support is therefore necessary before unseen trees can be scored at all. Full support is not, however, sufficient. Both CCD1+*ϵ* and KRegCCD cover every held-out tree, but KRegCCD consistently achieves higher predictive accuracy than CCD1+*ϵ*, with improvements ranging from 2.1 to more than 800 nats per tree. The difference reflects two aspects of KRegCCD. First, its split-expanded regCCD backbone represents more trees than the CCD1 backbone, leaving less for the escape mass to cover. Second, it allocates the escape probability among unsupported trees according to their clade structure, rather than uniformly as in CCD1+*ϵ*.

**Table 2:**
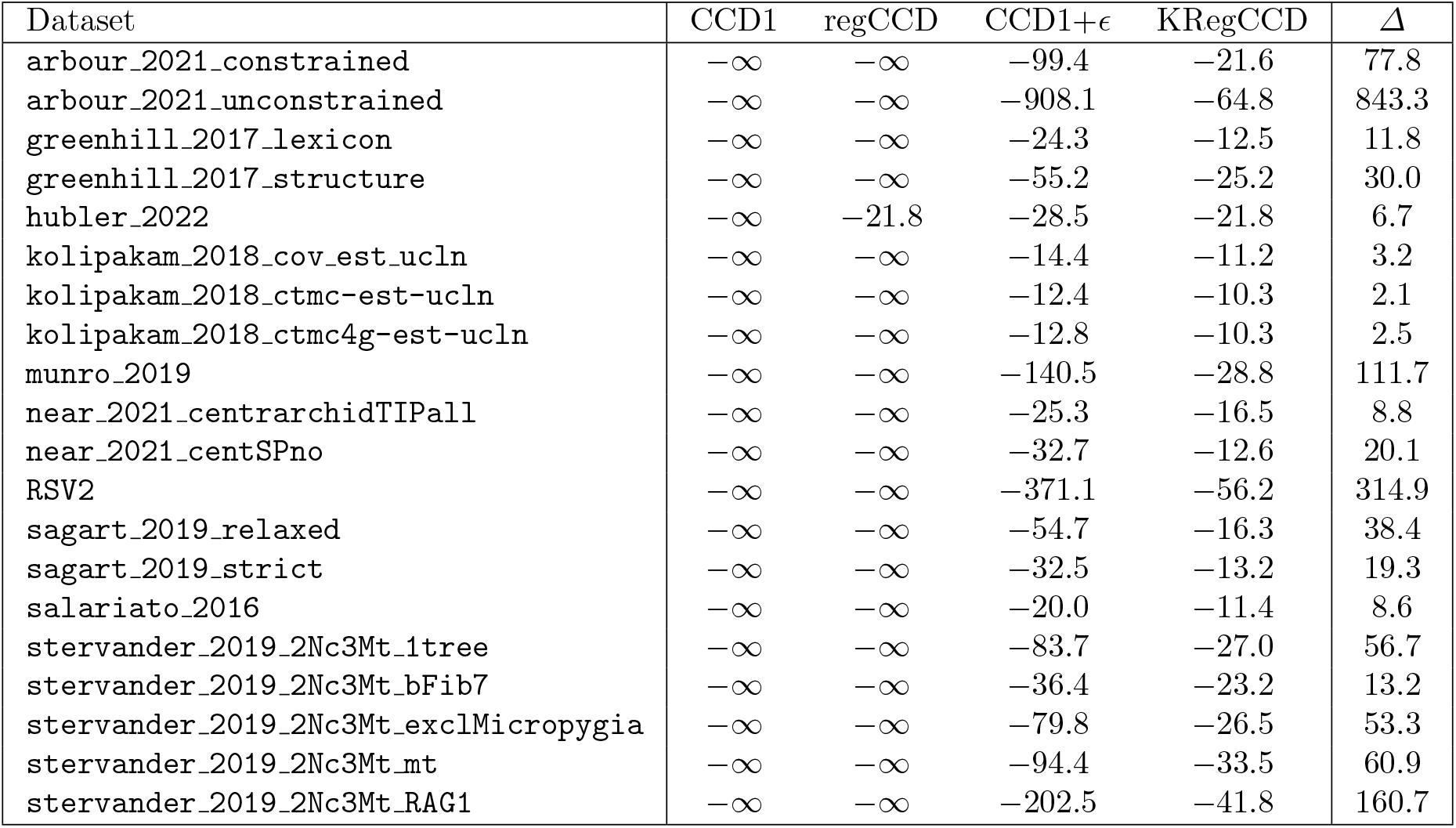
Mean held-out log probability per tree across empirical posterior tree sets with entropy ≥ 10. Models were trained on up to 1000 trees from the first half of each post-burn-in chain and tested on the same number from the second half (fewer for chains with fewer post-burn-in samples). *α* = 0.4; *μ* was fitted by cross-validation on the training half. *Δ* = KRegCCD − (CCD1 + *ϵ*) denotes the difference in mean held-out log probability.

**Fig. 9:**
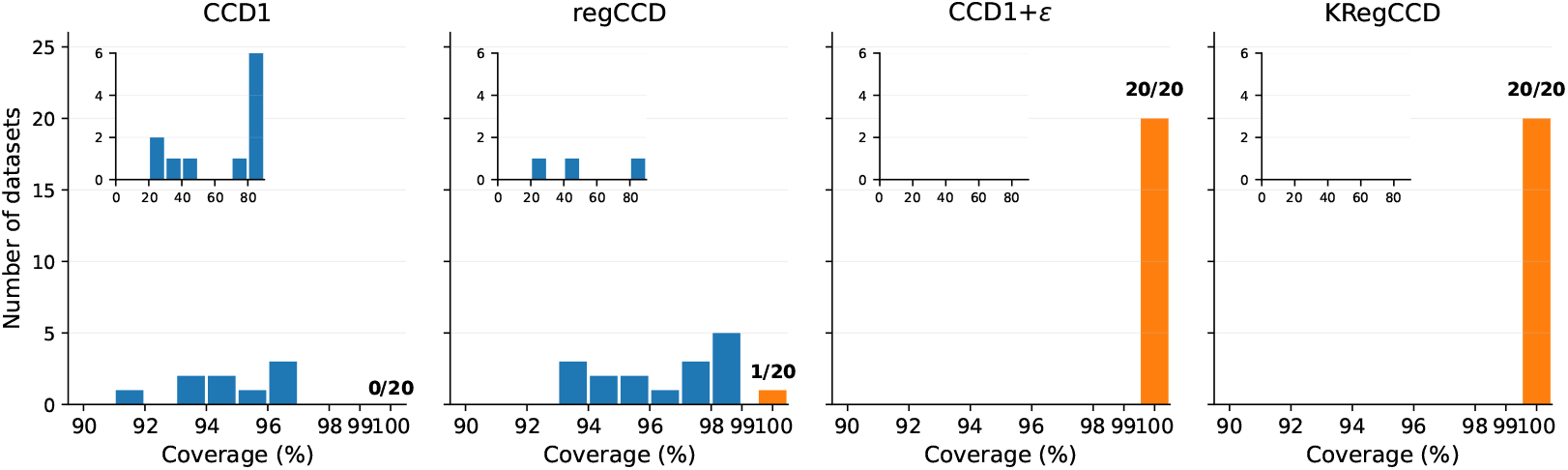
Held-out-tree coverage across the datasets in Table 2. Bars show the number of datasets in each coverage range. The insets show the number of datasets with coverage from 0 ∼90%, while the main plots provide a magnified view of the 90 ∼ 100% range. CCD1 and regCCD generally have incomplete coverage, whereas CCD1+*ϵ* and KRegCCD achieve 100% coverage for all datasets. **Alt text:** Bar charts compare held-out-tree coverage across real datasets for four models: CCD1, regCCD, CCD1+*ϵ*, and KRegCCD.

### 3.3 Point Estimate

We examined the effect of regularisation on point estimation by comparing the RF distances of the true trees to the MAP trees of different CCD models. Figure 10 shows the mean relative RF distance across different sample sizes for datasets `Yule[n]` and `Coal[n]`. Recall that the relative RF distance gives the percentage of the *n* − 2 clades in the true tree that are not correctly recovered by an estimator. For example, in Coal320, a relative RF distance of 0.16 means that about 51 of the 318 nontrivial clades differ from the true tree. While CCD0 remained the best-performing method, KRegCCD attained a slightly lower mean RF distance than CCD1 on seven of the eight datasets, the exception being `Coal40`, where the two are equal to within 0.01 clades. The margin grows with the size of the tree, from 0.08 clades on `Yule50` to 3.81 on Coal320. We also tested the pseudocount scheme *α* = prior (results provided in Appendix S2), which gave comparable performance to Fig. 10, and similarly outperformed CCD1.

**Fig. 10:**
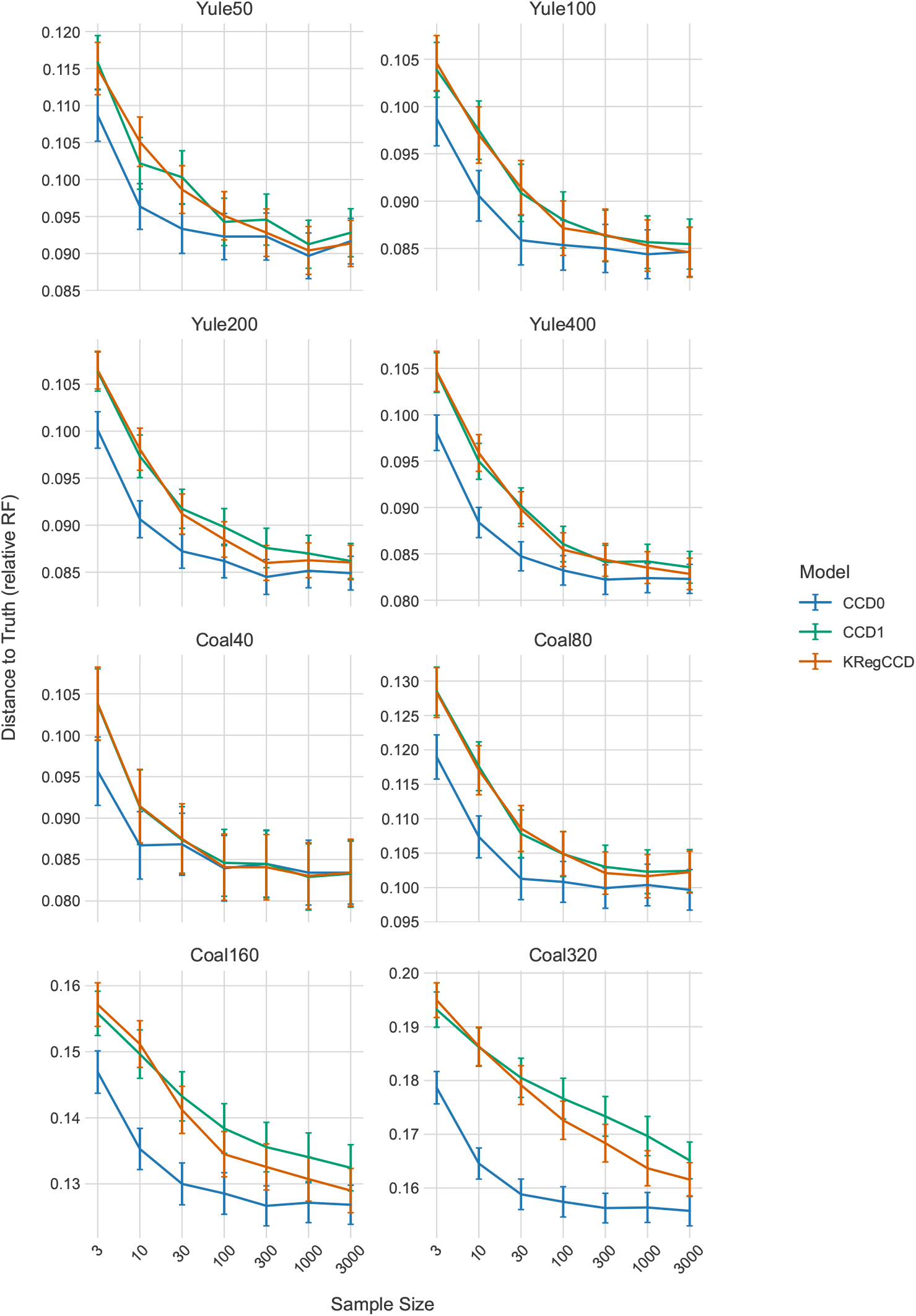
Accuracy of point estimates measured for models CCD0, CCD1, and KRegCCD with *α* = 0.4 and optimised *μ* in terms of the mean relative RF distance to the true tree for different sample sizes. CCD0 achieves the lowest mean RF distances, CCD1 the highest, while KRegCCD improves upon CCD1. **Alt text:** A comparison of mean relative RF distances across sample sizes shows differences in point estimation accuracy among CCD0, CCD1, and KRegCCD. (a) Optimal *α* values over 100 replicates for each dataset of `Yule[n]` and `Coal[n]`, coloured by the number of taxa, with three different subsample sizes (denoted as “sub”). The most common *α* is approximately 0.3 *∼* 0.4. As the subsample size increases, the optimal *α* decreases slightly. **Alt text:** Distributions of optimal *α* values across simulated datasets. (b) Optimal *α* distribution over fifty-nine RealData tree sets (excluding those with >5% unique-clade trees). Median *α* = 0.29, with one outlier at *α* = 3.75. **Alt text:** A distribution of optimal *α* values across real datasets.

One-sided paired Wilcoxon signed-rank tests over the 100 replicates (Table 3) find the improvement of KRegCCD over CCD1 significant only for Coal320 (*p* = 0.048), despite the favourable means. Reporting split-only regularisation (*α* = 0.5) alongside it shows why: split smoothing moves the MAP tree less often but far more reliably, beating CCD1 by 72 to 10 on Coal320 and 39 to 10 on Coal160, against 63 to 33 and 45 to 25 under clade regularisation. Escape mass redistributes probability near the mode as well as in the tail, so it perturbs the maximiser more readily, leaving the mean gain unchanged (104.10 against

**Table 3:** Statistical comparison of point estimation accuracy (subsample size = 1000). Mean RF distances to the true tree over 100 replicates, with one-sided paired Wilcoxon signed-rank tests. W/L/T gives the number of replicates in which the first model is closer to the true tree, further from it, and returns the identical tree; tests are computed on the non-tied replicates. *regCCD* denotes split-only regularisation (*α* = 0.5), included to separate the effect of split smoothing from that of clade regularisation. Significance: ^*^*p* < 0.05, ^**^*p* < 0.01, ^***^*p* < 0.001.

| Dataset | Mean RF Distance |  |  |  | KRegCCD vs CCD1 |  | regCCD vs CCD1 |  | CCD0 vs KRegCCD |  |
| --- | --- | --- | --- | --- | --- | --- | --- | --- | --- | --- |
| | CCD0 | KRegCCD | regCCD | CCD1 | $p$ | W/L/T | $p$ | W/L/T | $p$ | W/L/T |
| Yule50 | 8.6 | 8.7 | 8.7 | 8.8 | 0.300 | 28/24/48 | 0.049* | 4/0/96 | 0.586 | 31/32/37 |
| Yule100 | 16.5 | 16.7 | 16.7 | 16.8 | 0.507 | 28/27/45 | 0.062 | 6/2/92 | 0.319 | 39/35/26 |
| Yule200 | 33.7 | 34.2 | 34.2 | 34.5 | 0.410 | 30/27/43 | 0.007** | 15/4/81 | 0.160 | 51/39/10 |
| Yule400 | 65.6 | 66.5 | 66.4 | 67.0 | 0.321 | 36/31/33 | 9.1e-4*** | 20/8/72 | 0.085 | 51/38/11 |
| Coal40 | 6.3 | 6.3 | 6.3 | 6.3 | 0.572 | 21/25/54 | 0.681 | 1/1/98 | 0.476 | 31/29/40 |
| Coal80 | 15.7 | 15.9 | 15.9 | 16.0 | 0.304 | 32/26/42 | 0.038* | 6/2/92 | 0.322 | 43/36/21 |
| Coal160 | 40.2 | 41.3 | 41.2 | 42.4 | 0.149 | 45/25/30 | 2.5e-6*** | 39/10/51 | 0.048* | 60/33/7 |
| Coal320 | 99.5 | 104.1 | 104.2 | 107.9 | 0.048* | 63/33/4 | 3.2e-12*** | 72/10/18 | 0.037* | 58/39/3 |

104.18 on Coal320) but the paired comparison less consistent. Point estimation is therefore not where clade regularisation contributes: KRegCCD is comparable to CCD1 rather than reliably better, and CCD0 remains the most accurate throughout.

### 3.4 Regularisation Parameter Selection

To identify which regularisation parameters provide the most accurate posterior estimates, we optimised *α* and *μ* by maximising the mean log probability of held-out trees using cross-validation.

Fig. 11a shows the distribution of optimised *α* for regCCD under the supported tree-space for `Yule[n]` and `Coal[n]`. For the majority of datasets, *α* ≈ 0.3 ∼0.4 appears to be the most likely optimal value. Each dataset was tested with three subsample sizes (300, 1000, 3000), and we observe a slight decrease in the optimal *α* as the subsample size increases.

**Fig. 11:**
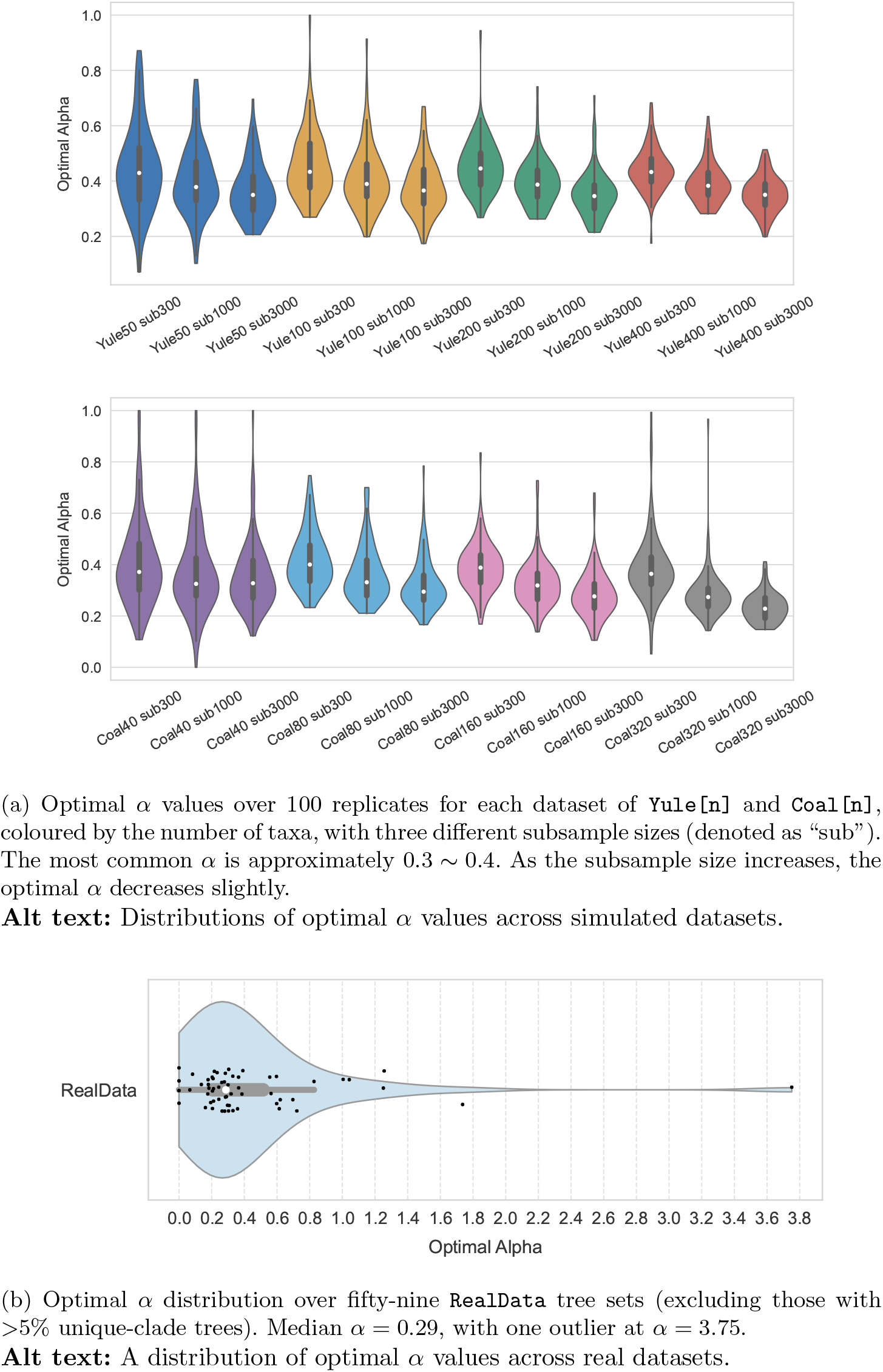
The distribution of optimal *α* for simulated and real datasets.

Fig. 11b shows the distribution of optimised *α* for regCCD under the supported tree-space for `RealData`, excluding tree sets where more than 5% of trees contain unique clades (see Table S1). While optimal values span a broad range, the density is highest around *α* ≈ 0.2 ∼ 0.4, with a median of 0.29. Most values fall below *α* = 1 (standard Laplace smoothing), with one outlier at *α* = 3.75. Six tree sets have optimal *α* ≥ 1 (Table 4); the differences in mean log probability between optimal and median *α* are small for all of them, suggesting relatively flat posteriors where regularisation performs similarly across *α* values.

**Table 4:** Tree sets with optimal *α* ≥ 1 and the difference in mean log probability between optimum and median.

| Dataset | Optimal $\alpha$ | $\Delta$ Mean Log(P) |
| --- | --- | --- |
| brandrud_orchid [12] | 1.26 | 1.77E-5 |
| foote_orca (subsample1) [23] | 1.74 | 0.0054 |
| hubler_language [30] | 3.75 | 0.0012 |
| mathers_aphids [45] | 1.04 | 1.68E-5 |
| mays_avian (codonpartitioned) [46] | 1.01 | 5.02E-7 |
| stange_catfishes (ariidae) [65] | 1.25 | 2.89E-5 |

Fig. 12 shows the distribution of optimised *μ* for KRegCCD for Yule[n] and Coal[n] with subsample sizes 300, 1000 and 3000. We observe that the optimal *μ* decreases as the subsample size increases. For Coal[n], the optimal *μ* increases with the number of taxa.

**Fig. 12:**
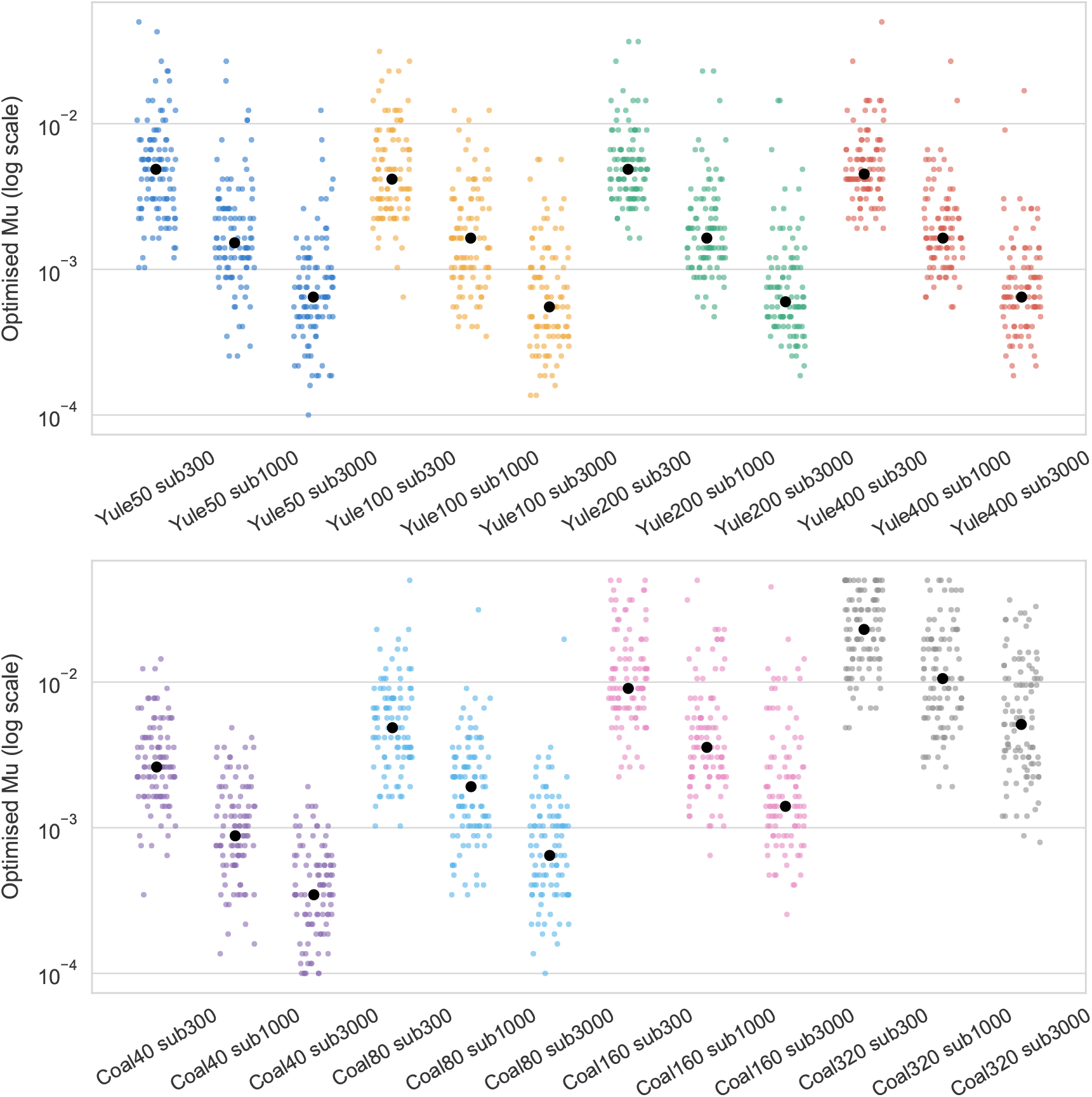
Optimal *μ* values over 100 replicates for each dataset of Yule[n] and Coal[n], coloured by the number of taxa, with three different subsample sizes (denoted as “sub”). As the subsample size increases, the optimal *α* decreases slightly. **Alt text:** Distributions of optimal *α* values across simulated datasets.

### 3.5 Bayesian Tree Topology Testing

Finally, we demonstrate an application enabled by KRegCCD: topology testing with Bayesian posterior tree distributions. Because KRegCCD provides a probability distribution over the space of all possible topologies, it allows arbitrary candidate trees to be evaluated against a posterior distribution. For example, a maximum-likelihood tree can be tested to determine whether it is supported by a Bayesian posterior, summary trees from independent Bayesian analyses can be compared to assess whether they are mutually supported, and posterior tree distributions obtained under different evolutionary models or datasets can be compared to determine whether they imply statistically different topological hypotheses. Specifically, we evaluate whether a candidate tree falls within the 95% credible set of a posterior distribution; exclusion from this credible set indicates that the candidate topology is not supported by that posterior.

For each of the datasets `Percidae, Liocichla` and `Rallidae`, we conducted pairwise comparisons of its posterior tree sets by computing the credible level of the MAP tree from one tree set in the CCD of the other. We used CCD0 for MAP trees and KRegCCD for building credible sets. The results are shown in Table 5. Each table entry gives the credible level of the point estimate (row) in the credible set of another distribution (column), where 0 denotes the highest level and 1 the outermost. Entries with values > 0.95 (in the tail of the distribution) indicate incompatible distributions and are highlighted in bold orange.

**Table 5:** Posterior tree distribution comparison across different models. Each cell shows the credible level of the MAP tree (row) within the credible set of the posterior distribution (column). Values outside the 95% credible set are highlighted in bold orange.

|  | constrained | unconstrained |
| --- | --- | --- |
| constrained |  | **1** |
| unconstrained | **1** |  |

(b) *Liocichla*: codon partitioned vs gene partitioned
|  | codon partitioned | gene partitioned |
| --- | --- | --- |
| codon partitioned |  | 0.01 |
| gene partitioned | **0.99** |  |

(c) *Rallidae*: combined five-marker shared-tree analysis vs mitochondrial-only tree analysis.
|  | 2Nc3Mt | 2Nc3Mt mt |
| --- | --- | --- |
| 2Nc3Mt |  | **1** |
| 2Nc3Mt mt | **1** |  |

Across the three datasets, we observed that posterior tree distributions inferred under different models can show substantial topological disagreement. In Percidae and Rallidae, the posterior distributions were mutually incompatible, with each analysis rejecting the MAP tree of the other. In Liocichla, the disagreement was asymmetric: the codon-partitioned MAP tree was supported by the gene-partitioned posterior, whereas the gene-partitioned MAP tree was not supported by the codon-partitioned posterior. Additional posterior comparisons between a larger set of evolutionary models across more datasets are provided in Supplementary Material Appendix S4.

## 4 Discussion

We introduced KRegCCD, a novel family of full-support models that improves posterior estimation of phylogenetic trees. KRegCCD can be constructed efficiently from a set of posterior trees. We compared the performance of our model with two alternative CCD models, and developed methods to optimise the regularisation parameters for a given tree set. Additionally, we showed its utility for Bayesian tree topology testing, providing a new framework for evaluating candidate topologies against posterior tree distributions.

Our experiments demonstrated that KRegCCD outperforms CCD0 and CCD1 in capturing the structure of the posterior distribution. CCD0 substantially underestimated posterior support and produced an overly loose fit of the posterior, consistent with the findings of Berling et al. [5]. The performance of CCD0 also showed little improvement as the number of samples increases, indicating again its limitation. CCD1 captured the structure of the posterior distribution better than CCD0 across all sample sizes, but it still underestimated posterior support.

This is a bias–variance trade-off in the estimation of the tree distribution. Write the expected discrepancy between a fitted CCD and the true posterior as the sum of an approximation error, the closest the model class can come to the posterior given unlimited samples, and an estimation error, the fluctuation of the fitted model across replicate MCMC samples; call these bias and variance. CCD0 has one free parameter per observed clade, from which its clade split probabilities are derived, whereas CCD1 has one per observed clade split; a clade generally admits several splits, so CCD1 has the larger parameter count, even though the CCD0 graph has more edges, its extra edges being determined by the clade parameters rather than estimated independently. Each CCD0 parameter is estimated from all *N* sampled trees, while a CCD1 conditional probability is estimated from only the *f* (*C*) trees containing its parent clade, which for rare clades is a handful. CCD0 therefore has the lower variance and the higher bias, and CCD1 the reverse. The sample-size trends bear this out: CCD0 improves little as more trees are added, an approximation error that does not vanish with sample size, whereas CCD1 improves steadily. This also explains why the two rank oppositely on our two criteria: low variance makes CCD0’s mode stable across samples, giving the most accurate point estimate, while its high bias gives the worst calibrated distribution. Smoothing is the standard remedy, trading a little bias for a larger reduction in variance in the low-bias model, and this is what KRegCCD does. KRegCCD demonstrated strong performance overall, with accuracy improving as the sample size increased.

The optimisation of *α* and *μ* for Yule[n] and Coal[n] revealed a mild trend in which smaller values of both parameters tended to provide better estimates as the sample size increased. This suggests that, as more trees become available, and thus more clades and clade splits are present in the CCD, less regularisation is required to achieve accurate posterior estimates. With increasing sample size, the observed tree set provides better coverage of the tree space, reducing the need to reserve probability mass for unexplored regions. The large variance in optimal *α* and *μ* across tree sets indicates that it is worthwhile to optimise them for each tree set individually, rather than relying on a fixed value.

Among the included `RealData`, six tree sets have optimal *α* ≥ 1 (standard Laplace smoothing or above). For all of these, the differences in mean log probability between optimal and median *α* are small (Table 4), suggesting relatively flat posteriors where smoothing performs similarly across *α* values.

For split regularisation to be meaningful, the posterior tree set must contain enough clade information. Split regularisation works by smoothing the probabilities of clade splits. If the posterior does not provide reliable estimates of clades — for instance, when each sampled tree introduces new clades — then the corresponding split probabilities will also be poorly estimated. We recommend excluding datasets where more than 5% of trees contain unique clades, as this indicates that clade space is not yet well-sampled, even if tree space is not expected to be.

Regularisation also assumes that an absent clade is absent because sampling was incomplete. Where it is instead excluded by the signal itself, a zero is the correct estimate and smoothing is harmful. Posteriors that are multimodal in topology, as arise under recombination, hybridisation or horizontal gene transfer [19, 73], are the clearest case: with *AB* ∥ *CD* and *AC* ∥ *BD* both well supported and *AD* ∥ *BC* genuinely unsupported, CCD1 represents the exclusion correctly while CCD0 and KRegCCD assign *AD* ∥ *BC* probability it does not merit, and CCD1 will be the better model. Fitting *μ* by cross-validation makes the cost visible rather than hidden, since a multimodal posterior should drive the fitted escape mass down, but it does not remove it.

A related question is how to assess how well a CCD approximates the posterior it is built from. Because the posterior probability of an individual topology cannot be estimated from the sample, the natural target is not the full-topology distribution but its lower-dimensional marginals. Clade and clade-split frequencies are the coarsest such marginals and are reliably estimated, but they constrain the distribution only weakly. Intermediate statistics, such as the induced distribution over the topologies of three- or four-taxon subtrees, interpolate between single-clade marginals and the full topology: they remain low-dimensional enough to be estimated accurately by Monte Carlo, yet capture higher-order structure that clade frequencies alone do not. Comparing the subtree-topology distributions implied by a CCD against those estimated directly from the posterior sample therefore provides a goodness-of-fit assessment at a controllable resolution, and offers a principled way to detect where a model such as CCD0 or KRegCCD places probability mass that the posterior does not support. We regard this as a promising direction for validating tree-distribution models.

With respect to point estimates, KRegCCD slightly improves upon CCD1, but does not reach the accuracy of CCD0. KRegCCD should therefore be preferred over CCD1 for both point estimation and capturing the overall structure of the posterior distribution, though CCD0 remains the preferred choice for point estimates. Given a sufficiently large number of uncorrelated samples, the CCD1 and thus KRegCCD MAP tree are expected to perform comparably to the CCD0 MAP tree.

KRegCCD improves the base model along a different axis from models that enrich the conditional structure, such as the subsplit Bayesian networks of Zhang and Matsen [77]. Such models increase the conditional order, letting each split depend on a larger context than its immediate parent clade, and so capture higher-order dependencies in the posterior. KRegCCD instead retains the firstorder conditional structure of CCD1 and addresses a separate problem, namely the support of the model. CCD1 fails not because its dependency structure is too simple, but because finite sampling leaves parts of tree space unsupported, so that it assigns zero probability to unobserved clades and splits. Smoothing fills that support, which is the appropriate remedy for a coverage limitation rather than a dependency limitation. The two directions are complementary: a richer conditional model could be regularised in the same way to give it full support. In terms of computational efficiency, KRegCCD has essentially the same construction time as CCD0, as both methods go through the same expansion step. The additional cost is computing the escape coefficients *N*_*i*_(*C*) of Eq. (4), which requires enumerating partitions of each clade *C* into observed subclades rather than simply traversing *C* (*G*). Only the two leading coefficients are needed, and both are obtained from a single pass over the disjoint pairs of observed subclades of *C*: *N*_1_(*C*) by testing whether the complement of a pair is itself an observed subclade, and *N*_2_(*C*) by matching each pair against those covering its complement. The cost is therefore quadratic in the number of observed subclades of *C*, rather than the cubic cost of enumerating four-part partitions directly. As the coefficients are computed once per CCD and cached, this is feasible for typical large data sets, but the cost grows super-linearly with the number of sampled clades.

Given the constructed CCDs *D*^(−1)^, …, *D*^(−*F*)^, optimisation of the regularisation parameters via cross-validation is linear in the number of taxa *n* and the number of sampled trees *N*. Neither parameter requires the CCD graph to be rebuilt: *α* enters only through the smoothed CCPs *θ*_*α*_(*S*), and *μ* only through *ϵ*(*C*), which is re-solved from the (*μ*-independent) coefficients *N*_*i*_(*C*). Each candidate value is therefore evaluated in a single traversal of each held-out tree. We recommend optimising *α* and *μ* via cross-validation whenever feasible.

The topology-testing framework presented here provides a practical way to compare phylogenetic hypotheses across analyses. Rather than relying on likelihood-based comparisons, candidate topologies can be evaluated directly against posterior tree distributions using their credible levels. Although we demonstrated this approach using CCD-MAP trees from posterior distributions inferred under alternative evolutionary models, the framework is applicable to any candidate tree defined on the same taxon set, including trees inferred by maximum likelihood, proposed in previous studies, or representing specific evolutionary hypotheses. This provides a new perspective to traditional model-comparison approaches by assessing whether different analyses produce statistically compatible posterior distributions.

## 5 Conclusion

In this study, we introduced KRegCCD, a family of full-support models for improving posterior estimation of phylogenetic trees. By combining observed clade and split information with a structured modelling of the unexplored regions of tree space, KRegCCD provides more accurate estimates of posterior tree distributions than existing CCD models. We showed that KRegCCD better captures posterior structure than both CCD0 and CCD1, while achieving point estimates comparable or better than CCD1. We also developed an efficient cross-validation approach for optimising the regularisation parameters for individual tree sets. Furthermore, we demonstrated that KRegCCD enables Bayesian tree topology testing, allowing arbitrary candidate topologies to be evaluated against posterior tree distributions through their credible levels. This provides a new framework for assessing the support and compatibility of phylogenetic hypotheses across different analyses.

A prerequisite for applying regularisation is that clade space is well-sampled; we recommend excluding datasets where more than 5% of trees contain unique clades. Future work could apply Chao estimators for species richness [14] to estimate the number of unseen clades and further refine criteria for assessing whether posterior samples are suitable for CCD-based analysis.

## Software Availability

The KRegCCD implementation is available in the CCD package at https://github.com/CompEvol/CCD.

## Data Availability

Data and code supporting this study are available from: https://doi.org/10.5281/zenodo.23075034.

## Supplementary Material

### S1 Further Results on Tree Distribution

For completeness, we provide here the accuracy of the estimate of the full posterior distribution for all the Yule[n] and Coal[n] datasets. The results are shown in Figs. S1 to S7.

Each plot shows, for a given dataset, the fraction of the testing trees in a range of CCD credible sets built from the training set and annotated with average L1 norms. The minimum L1 value for each sample size is highlighted in orange. Models CCD0, CCD1, and KRegCCD with *α* = 0.4 and optimised *μ* are compared across sample sizes of 100, 300, and 1000. CCD0 performs the worst and largely underestimates posterior support, CCD1 provides better estimates than CCD0, and KRegCCD consistently achieves the highest accuracy.

**Fig. S1:**
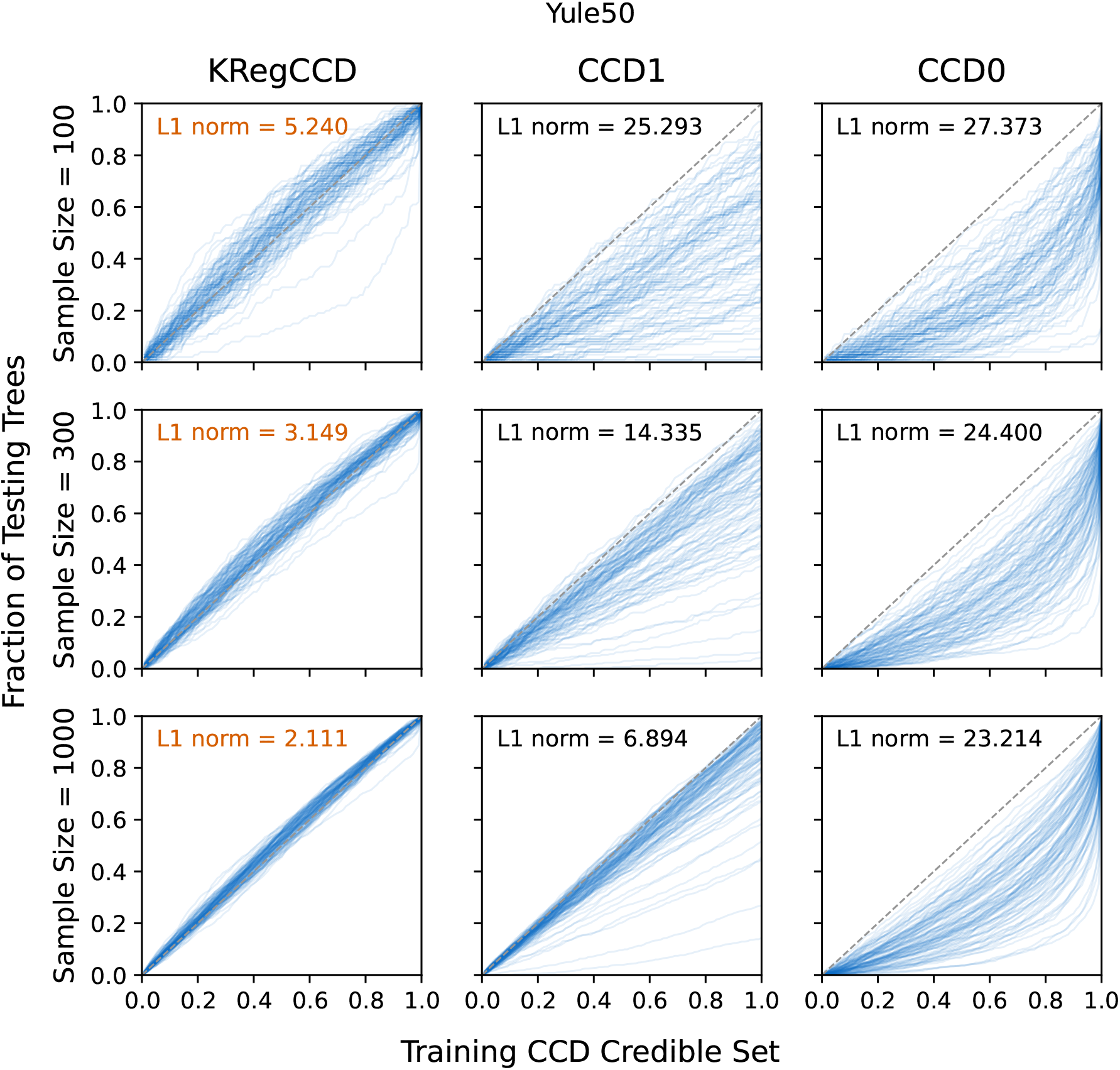
Posterior estimation accuracy for the Yule50 dataset. **Alt text:** A plot compares posterior estimation accuracy across different CCD models for Yule50.

**Fig. S2:**
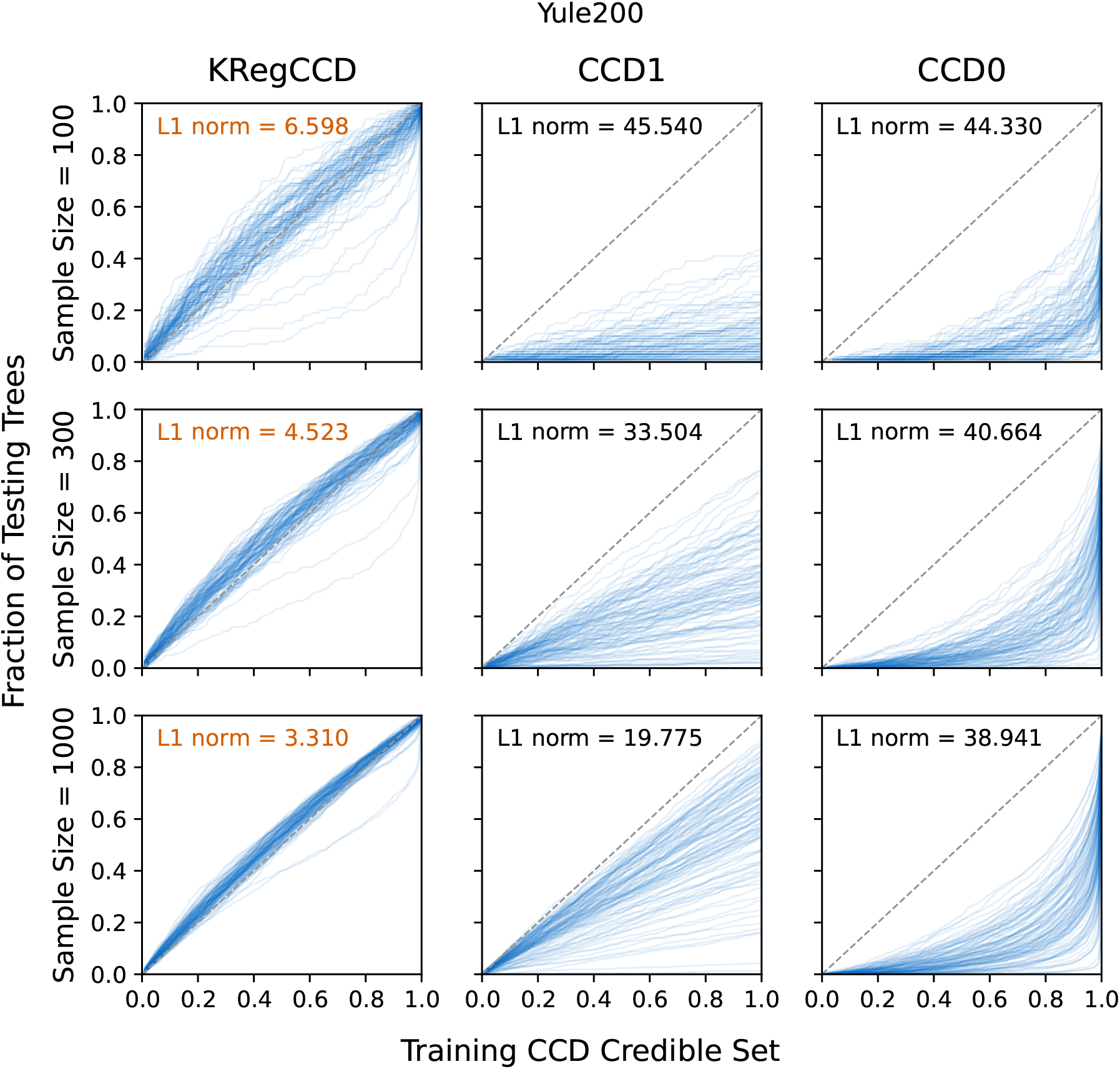
Posterior estimation accuracy for the Yule200 dataset. **Alt text:** A plot compares posterior estimation accuracy across different CCD models for `Yule200`.

**Fig. S3:**
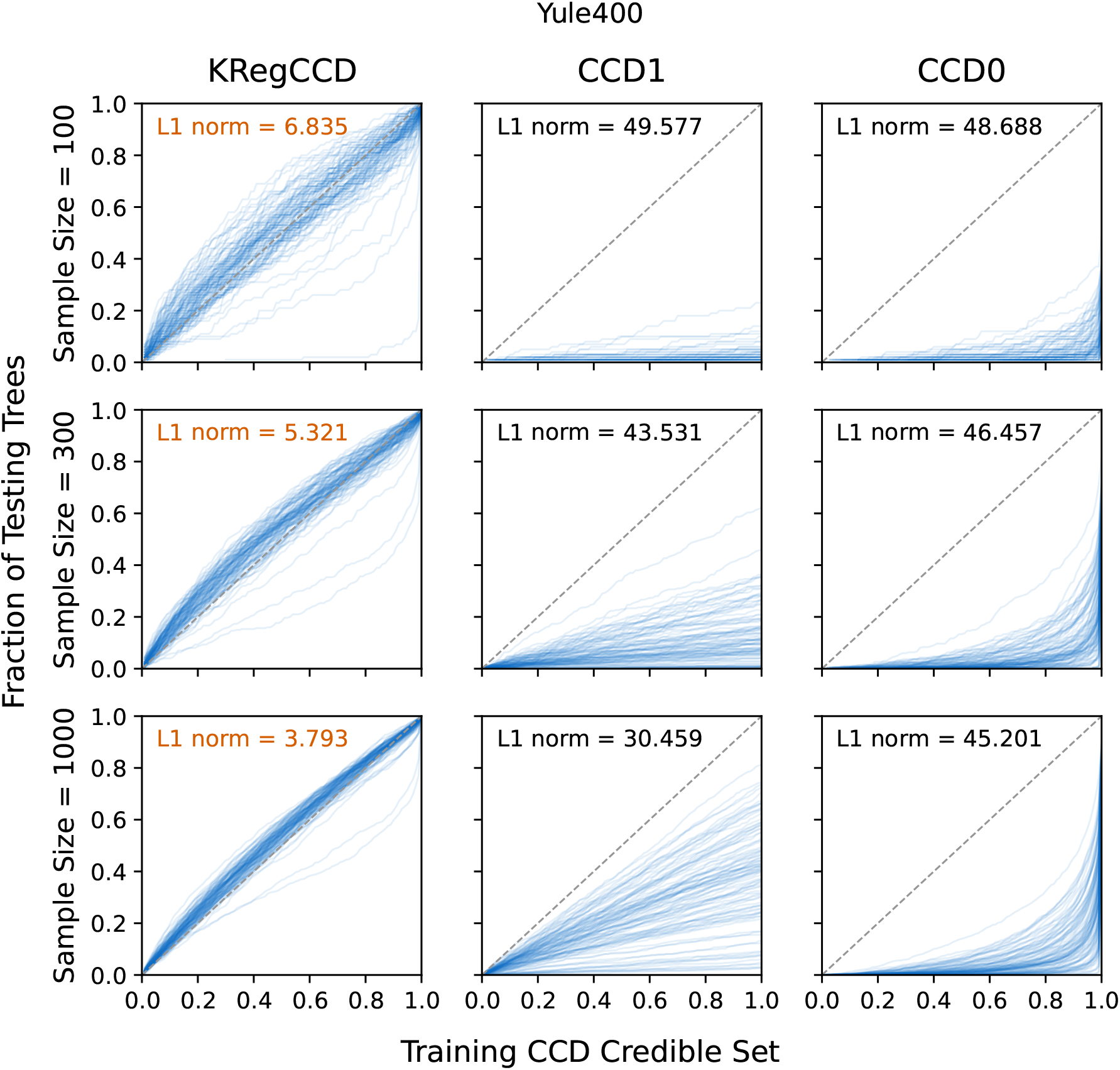
Posterior estimation accuracy for the Yule400 dataset. **Alt text:** A plot compares posterior estimation accuracy across different CCD models for `Yule400`.

**Fig. S4:**
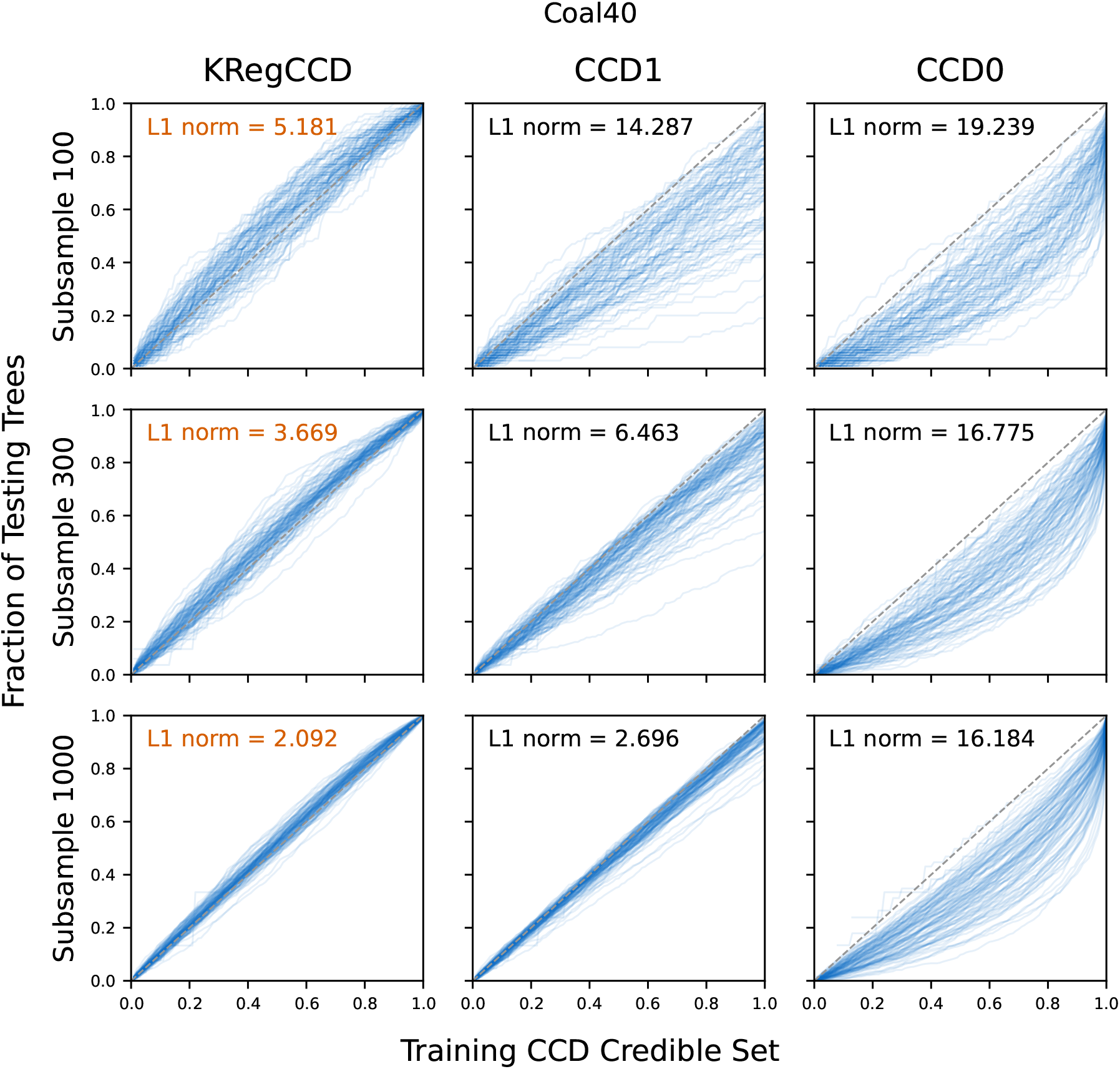
Posterior estimation accuracy for the `Coal40` dataset. **Alt text:** A plot compares posterior estimation accuracy across different CCD models for `Coal40`.

**Fig. S5:**
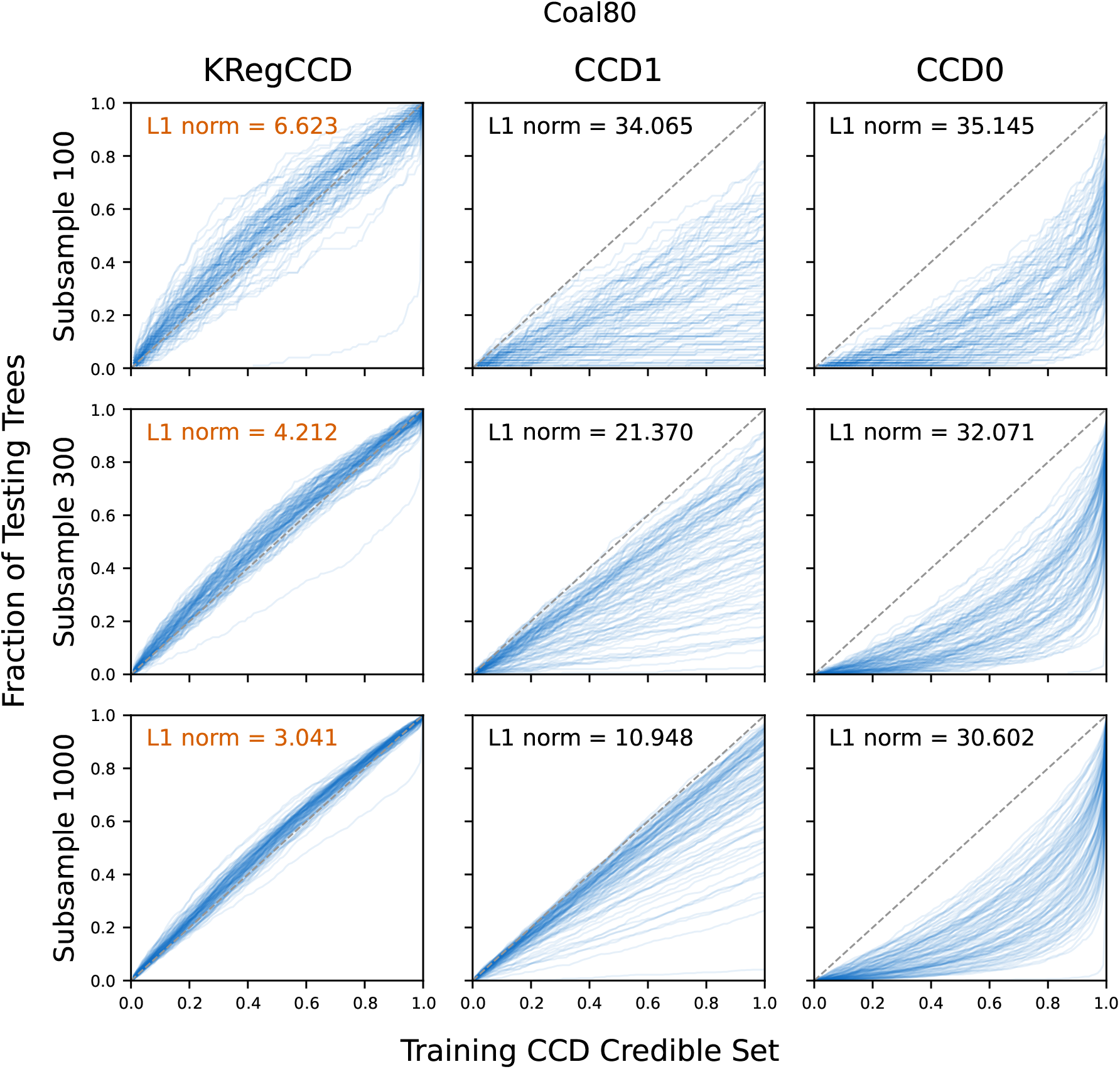
Posterior estimation accuracy for the `Coal80` dataset. **Alt text:** A plot compares posterior estimation accuracy across different CCD models for `Coal80`.

**Fig. S6:**
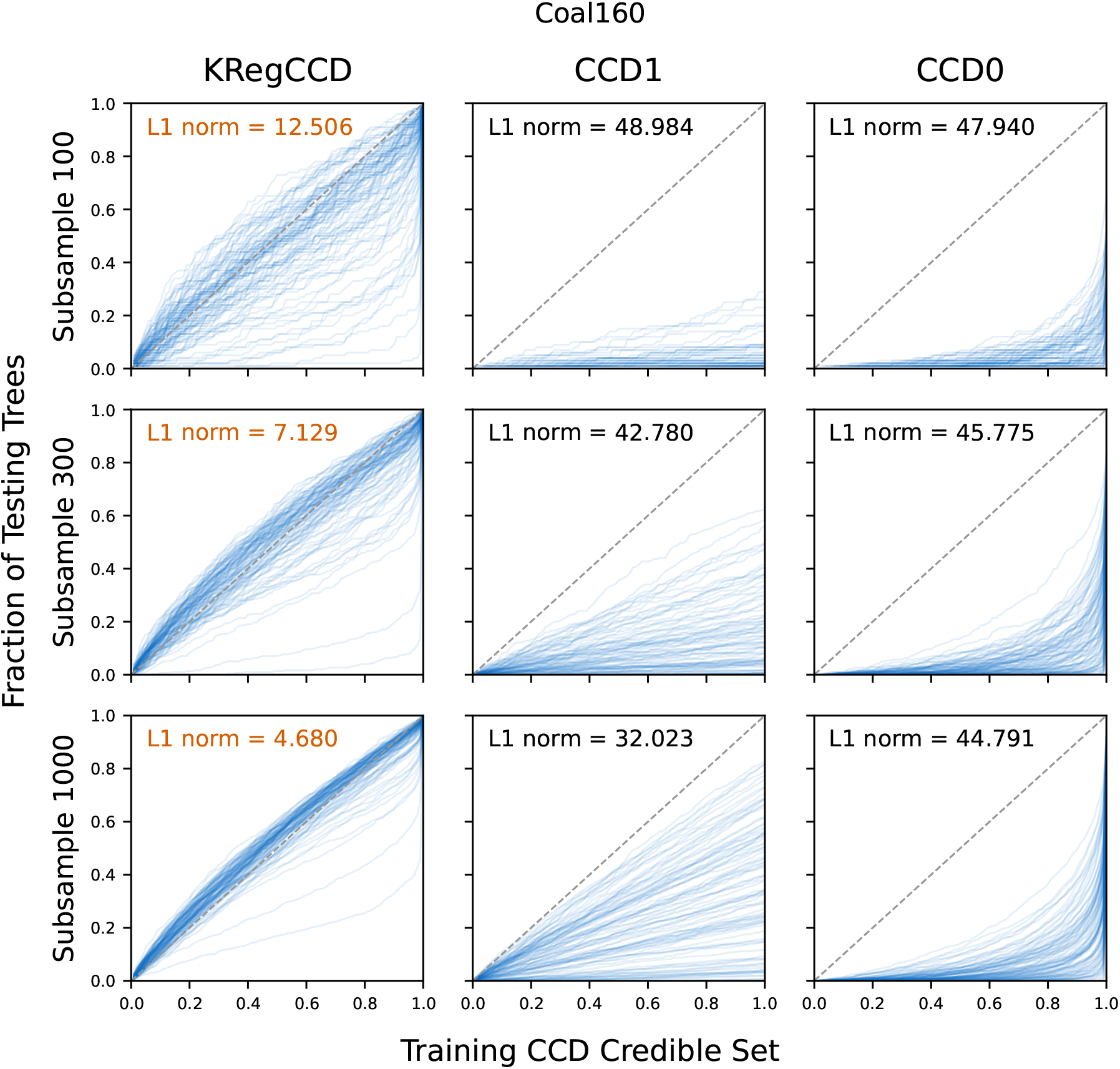
Posterior estimation accuracy for the `Coal160` dataset. **Alt text:** A plot compares posterior estimation accuracy across different CCD models for `Coal160`.

**Fig. S7:**
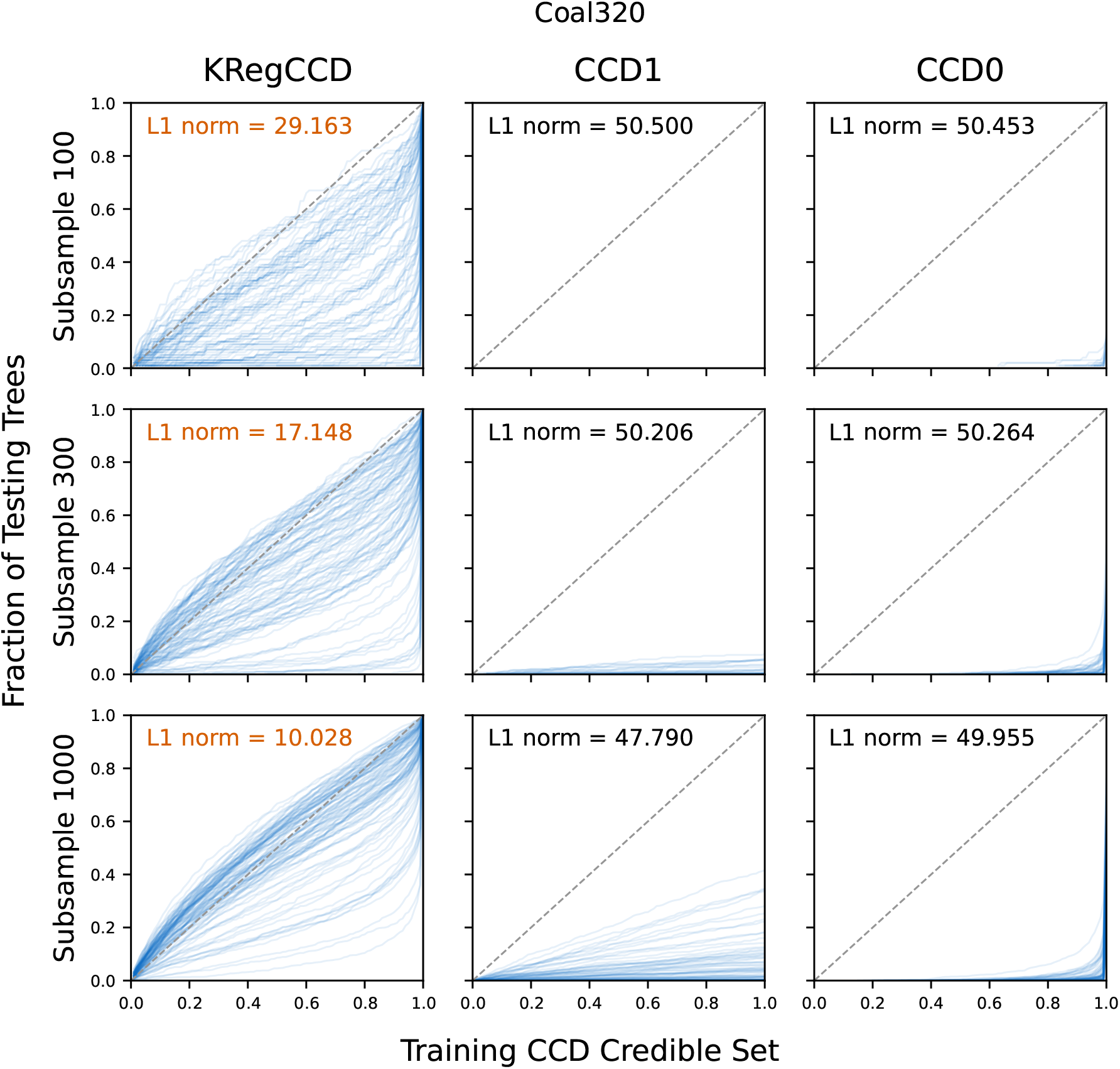
Posterior estimation accuracy for the `Coal320` dataset. **Alt text:** A plot compares posterior estimation accuracy across different CCD models for `Coal320`. The accuracy of the point estimates for the `Yule[n]` datasets with *α* = prior is shown in Fig. S8. regCCD (*α* = prior) produced results similarly to other smoothing variants, improving upon the point estimates obtained with CCD1, while CCD0 still provided the most accurate point estimates.

### S2 Further Results on Point Estimate

The accuracy of the point estimates for the `Yule[n]` datasets with *α* = prior is shown in Fig. S8. regCCD (*α* = prior) produced results similarly to other smoothing variants, improving upon the point estimates obtained with CCD1, while CCD0 still provided the most accurate point estimates.

**Fig. S8:**
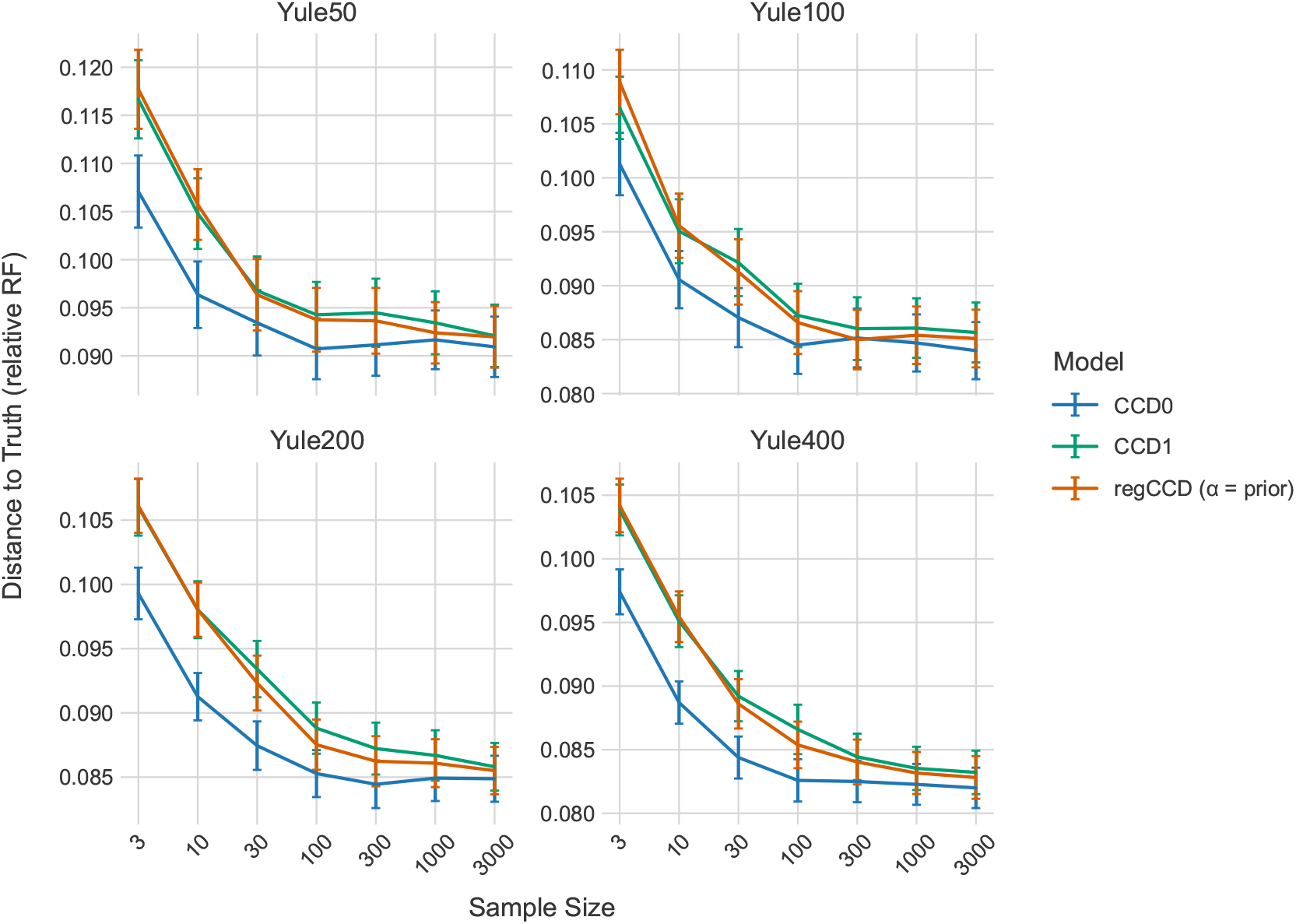
Accuracy of point estimates measured for models CCD0, CCD1, and regCCD with *α* = prior in terms of the mean relative RF distance to the true tree for different sample sizes. CCD0 achieves the lowest mean RF distances, CCD1 the highest, while regCCD improves upon CCD1. **Alt text:** A comparison of mean relative RF distances across sample sizes shows differences in point estimation accuracy among CCD0, CCD1, and regCCD with *α* = prior.

### S3 Real Data Summary

Table S1 lists all `RealData` tree sets used in this study, along with their CCD1 entropy, number of posterior samples, and the fraction of trees containing unique clades (which receive zero probability under leave-one-tree-out cross-validation). Tree sets where more than 5% of trees contain unique clades (highlighted in bold) indicate that clade space is not yet well-sampled, and regularisation results should be interpreted with caution.

**Table S1:**
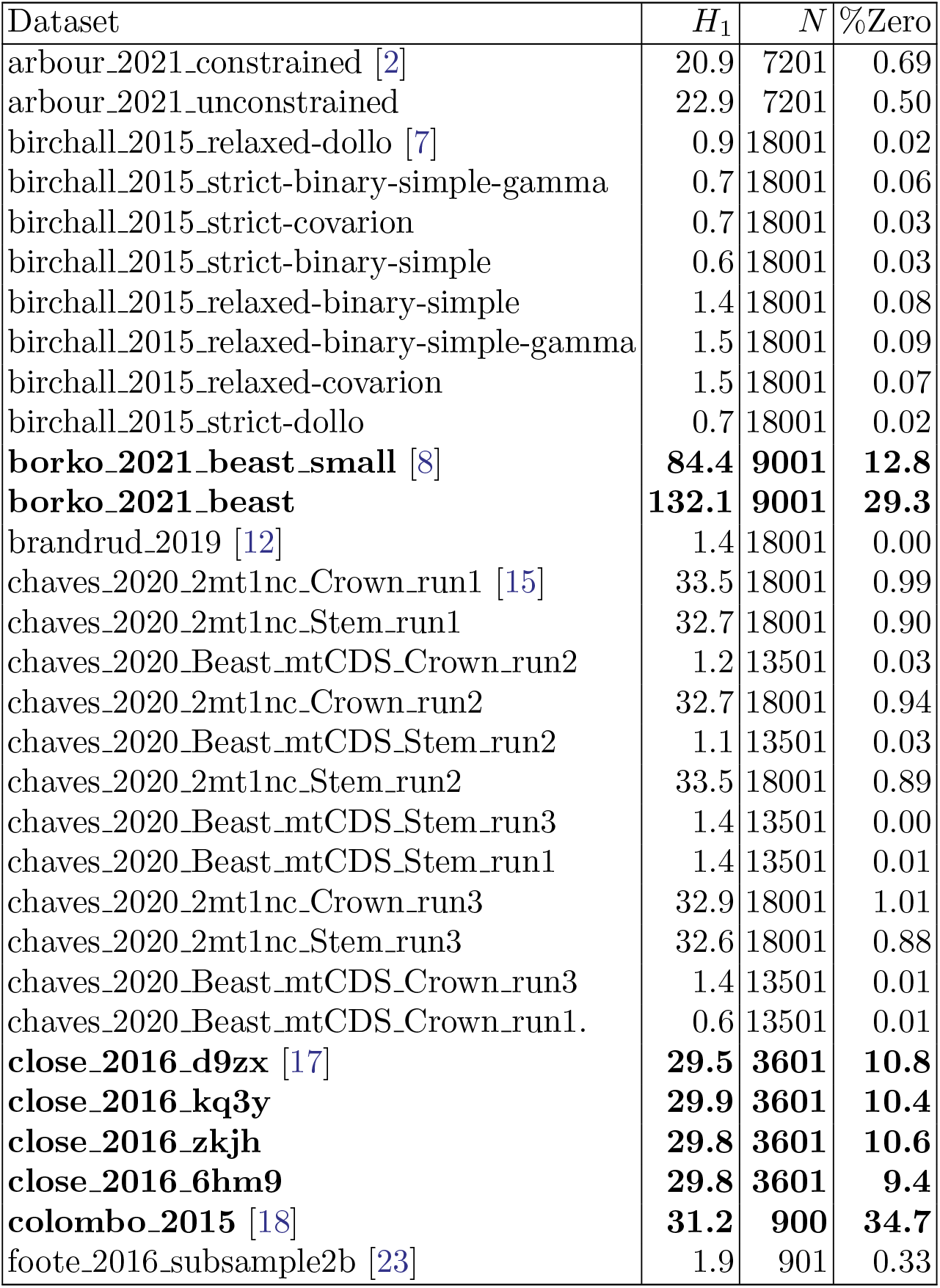
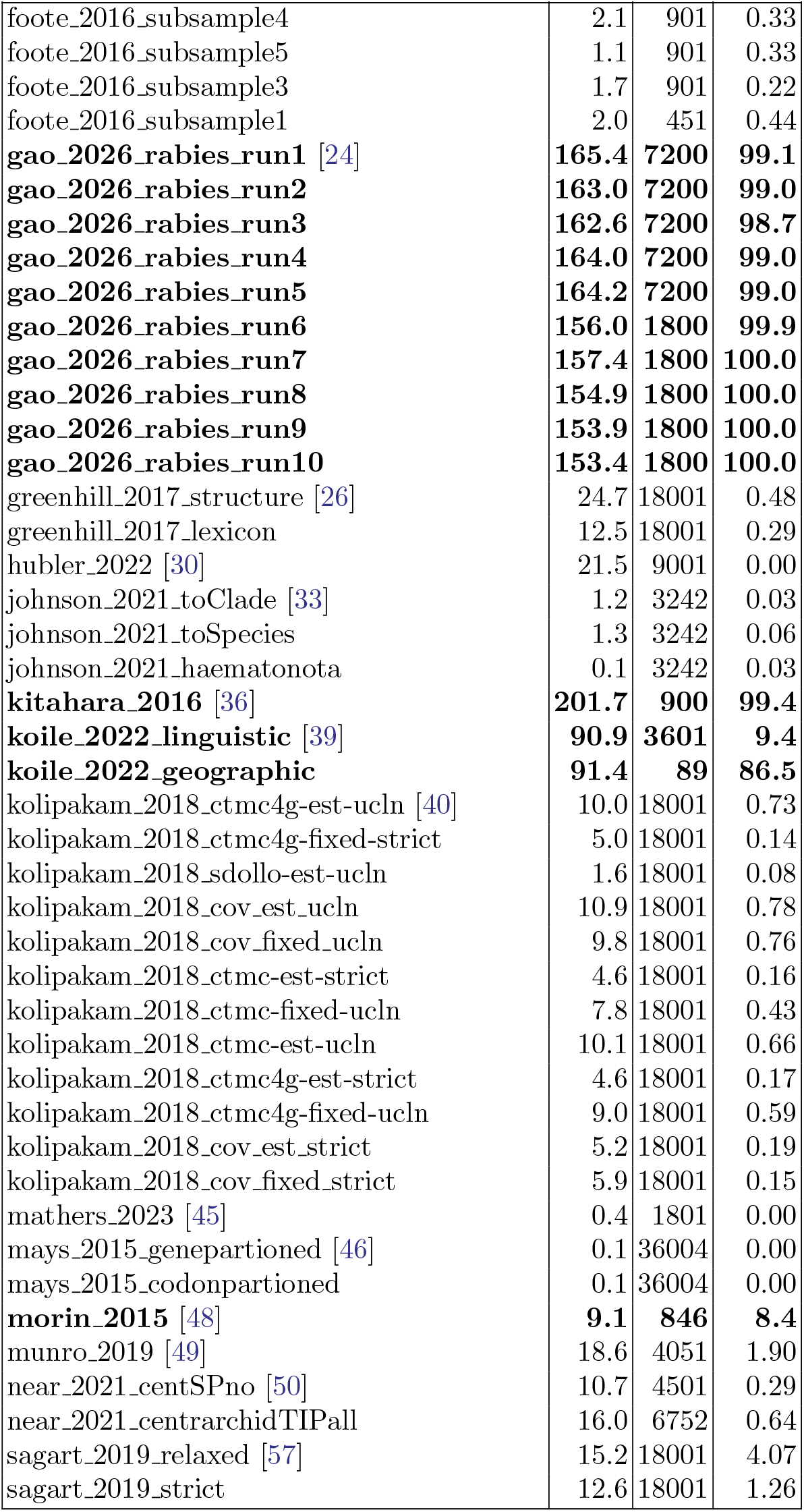
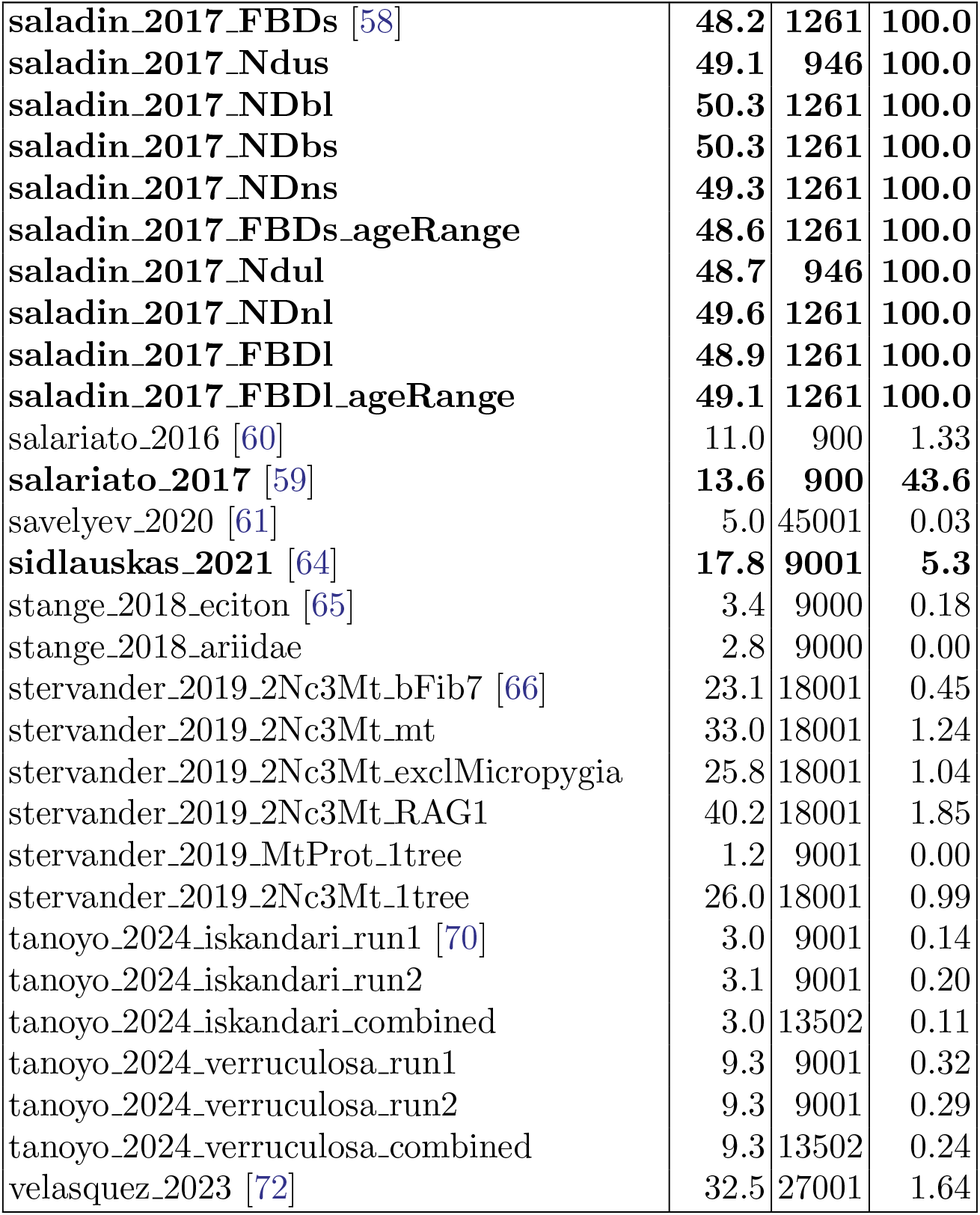
Summary of RealData tree sets. *H*_1_ denotes CCD1 entropy (nats), *N* is the number of posterior samples, and %Zero is the percentage of trees containing unique clades. Tree sets exceeding the 5% threshold are shown in bold. Studies contributing multiple tree sets are cited only at their first occurrence.

### S4 Further Results on Topology Testing

We further analysed several language datasets in `RealData` using regCCD to assess the compatibility of their posterior distributions:

`AUSN` Two sets of 20,000 trees on 81 Austronesian languages, one with lexical data and the other with grammatical data [26].

`CHAP` Eight sets of 20,000 trees on 10 Chapacuran languages [7]. Four cognate models were compared: simple binary continuous time Markov chain (CTMC) [9, 20, 71], simple binary continuous time Markov chain with gamma rate heterogeneity in four rate categories (CTMC4g) [74], covarion [52], and Dollo [1, 51], each with both strict and relaxed clock models [22].

`DRAV` Twelve sets of 20,000 trees on 20 Dravidian languages [40]. Four cognate models were compared: CTMC, CTMC4g, covarion, and Dollo. Two clock models were fitted, strict clock and uncorrelated lognormal relaxed clock [22]. Each cognate model was further tested with mutation rate estimated versus fixed [13].

`SNTB` Two sets of 20,000 trees on 50 Sino-Tibetan languages [57], one with strict clock and the other with relaxed clock.

The results are shown in Table S2. Because regCCD does not provide full support, some MAP trees have zero probability under the other posterior, indicated by −1 in the tables. Such zero-probability cases do not occur with KRegCCD. We observe that, for `AUSN`, the two posteriors are incompatible, consistent with the literature indicating that lexical and grammatical data measure different linguistic dimensions [26]. For `CHAP`, the Dollo model results in distributions that differ from those of the other cognate models, with the credible levels of the MAP trees from all other models exceeding 95% for relaxed Dollo and approaching the threshold (∼ 0.92) for strict Dollo. For `DRAV`, the Dollo model likewise produces distributions distinct from the other models. For both `DRAV` and `SNTB`, there is a substantial difference between the relaxed and strict clock model.

**Table S2:**
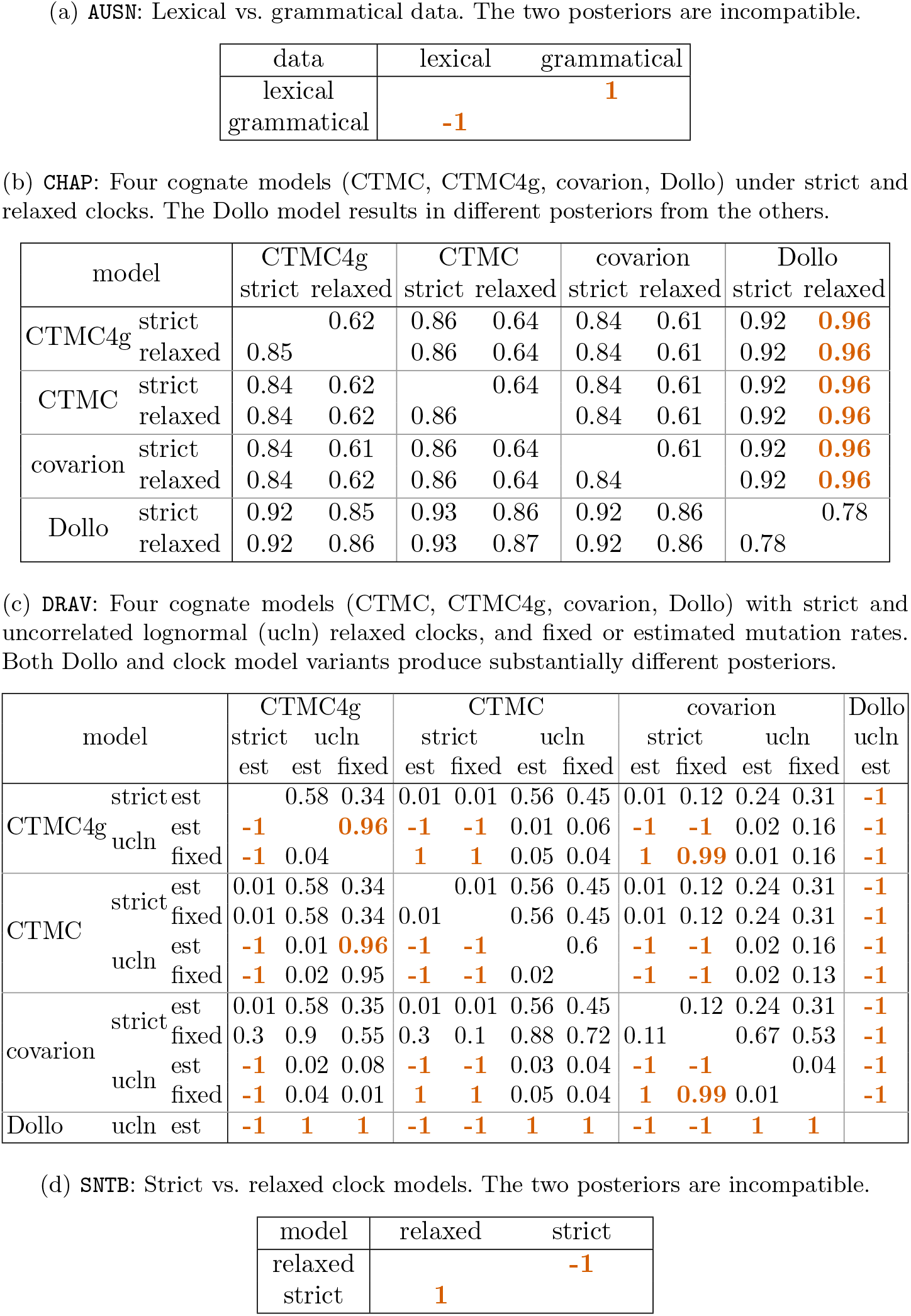
Posterior tree distribution comparison across data and models. Each cell shows the credible level of the MAP tree (row) within the credible set of the posterior distribution (column). Values outside the 95% credible set are highlighted bold orange; −1 indicates the MAP tree has zero probability under the other posterior (contains clades not observed in the other posterior tree sample).

